# Social Behavior Atlas: A computational framework for tracking and mapping 3D close interactions of free-moving animals

**DOI:** 10.1101/2023.03.05.531235

**Authors:** Yaning Han, Ke Chen, Yunke Wang, Wenhao Liu, Xiaojing Wang, Jiahui Liao, Yiting Huang, Chuanliang Han, Kang Huang, Jiajia Zhang, Shengyuan Cai, Zhouwei Wang, Yongji Wu, Gao Gao, Nan Wang, Jinxiu Li, Yangwangzi Song, Jing Li, Guodong Wang, Liping Wang, Yaping Zhang, Pengfei Wei

## Abstract

The study of social behaviors in animals is essential for understanding their survival and reproductive strategies. However, accurately tracking and analyzing the social interactions of free-moving animals has remained a challenge. Existing multi-animal pose estimation techniques suffer from drawbacks such as the need for extensive manual annotation and difficulty in discriminating between similar-looking animals in close social interactions. In this paper, we present the Social Behavior Atlas (SBeA), a novel computational framework that solves these challenges by employing a deep learning-based video instance segmentation model, 3D pose reconstruction, and unsupervised dynamic behavioral clustering. SBeA framework also involves a multi-camera setup to prevent occlusion, and a novel approach to identify individual animals in close social interactions. We demonstrate the effectiveness of SBeA in tracking and mapping the 3D close interactions of free-moving animals using the example of genetic mutant mice, birds, and dogs. Our results show that SBeA is capable of identifying subtle social interaction abnormalities, and the models and frameworks developed can be applied to a wide range of animal species. SBeA is a powerful tool for researchers in the fields of neuroscience and ecology to study animal social behaviors with a high degree of accuracy and reliability.

## Introduction

Close social interactions are critical for the survival and reproduction of animals^1^. However, the study of social behaviors has traditionally relied on rudimentary measures, such as the duration of time spent in specific areas during experiments like three-chamber tests^2^. To address this limitation, deep learning-based quantitative measurements have emerged as a potential solution^3^. In particular, there has been a surge of interest in developing multi-animal pose estimation and behavioral mapping techniques across various disciplines, including neuroscience and ecology^4^. Although single-animal pose estimation has been achieved with high accuracy through deep learning, accurately tracking and mapping the social behaviors of multiple animals remains a challenging task^5^.

Advanced multi-animal pose estimation toolboxes, such as Multi-animal DeepLabCut (maDLC) and Social LEAP Estimate Animal Poses (SLEAP), have enabled markerless and precise tracking of body parts for different species based on videography^6–8^. However, these techniques suffer from several limitations. Firstly, the high level of tracking precision necessitates a significant amount of manual annotation, which becomes increasingly laborious as the number of animals in the study increases. Secondly, occlusion can occur when multiple animals are present in the same video frame, resulting in poor inference about the behavior of each animal. Thirdly, in close social interactions between similar-looking animals^9^, it becomes challenging to distinguish between individual identities, particularly over extended periods of time^10^.

The Social Behavior Atlas (SBeA) offers a solution to the challenges proposed by existing multi-animal pose estimation techniques. Firstly, the number of manual annotations can be reduced by comprising two processes. The first is the acquisition of each animal’s contour. As fewer as 400 annotations are enough to separate adjacent animals. These data generate millions of labeled frames to train the deep learning-based video instance segmentation (VIS) model. Secondly, to address the issue of occlusion, SBeA employs multiple cameras to capture video streams, which are used to reconstruct 3D poses and prevent complete occlusion^11–13^. Thirdly, SBeA resolves the multi-animal identification problem by merging the contour of each animal with the characteristic identity of multiple view angles, which achieves over 90% identification precision without human data annotation. Furthermore, after solving those problems, inspired by the natural structures of social behavior, the unsupervised dynamic behavioral metric learning is finally designed. The behavioral metric is composed of a time-series low-dimensional representation of the behavior module. The behavioral mapping generates the social behavior atlas, and one-third of the cluster purity reaches over 95%. Using SBeA, we found the subtle social interaction abnormalities of Shank3B KO mice, which verifies the availability of SBeA. The models and frameworks developed for SBeA can be also applied to birds and dogs, showcasing its strong generalization abilities suitable for various application scenarios.

## Results

### SBeA: from multi-animal markerless 3D pose tracking to unsupervised social behavior mapping

The focus of SBeA is on quantifying the behavior of freely social animals comprehensively. It presents two significant challenges: pose tracking and behavior mapping. The pose tracking involves identifying the key body parts of each animal as well as their identities, which is particularly challenging when dealing with animals that look similar^10^. To address this issue, a novel free social behavior test paradigm has been developed that involves a multi-view camera array (Fig. 1a). This approach captures the animals covering more view angles and helps to overcome the challenge of frequent occlusion^11–14^. The camera array is used to capture images of a checkerboard for camera calibration, followed by videos of two free-moving animals for the social behavior test (Video capture phase 1, Fig. 1a). Finally, the array captures videos of single free-moving animals to facilitate animal identification without the need for human intervention (Video capture phase 2, Fig. 1a).

**Fig. 1.**
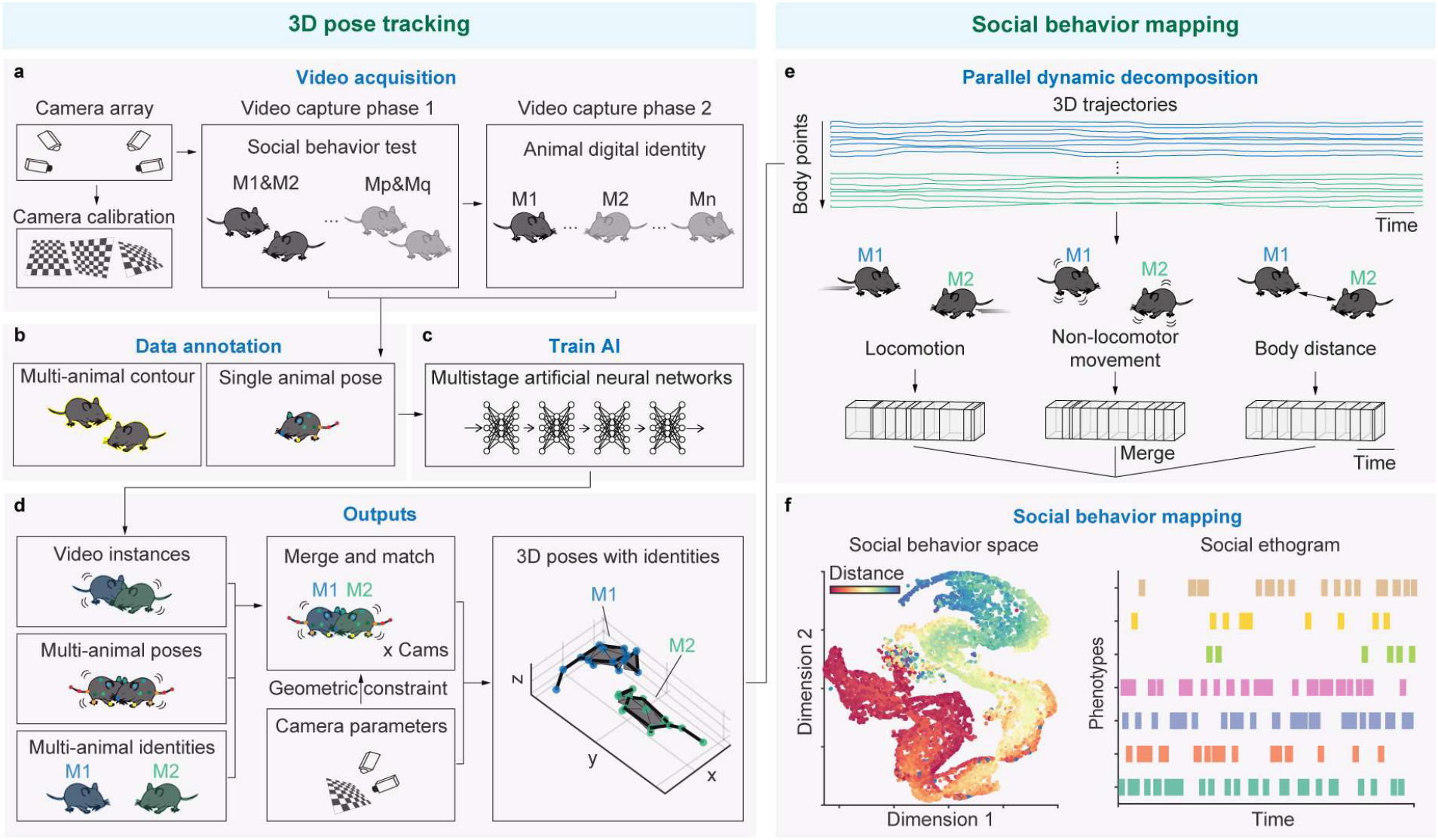
The architecture of Social Behavior Atlas. **a**, Video acquisition for free social behavior test. The camera array is used for behavioral capturing, and it is calibrated by checkboard images. There are two phase for beahvioral video capturing including social behavior test and animal digital identity. The phase 1 is to capture the videos of free-social interactions of two mice. The phase 2 is to capture the identities of each mice in phase 1. **b**, Data annotation for AI training. Social Behavior Atlas need the annotations of multi-animal contour and single animal pose. **c**, The multistage artificial neural networks for 3D pose tracking. **d**,The outputs of 3D pose tracking. Left: The outputs of AI including video instances, multi-animal poses, and multi-animal identities. Center: Combining video instances, multi-animal poses, and multi-animal identities with camera calibration parameters for 3D reconstruction with identities. Right: The visualization of 3D poses with identities. **e**, Parallel dynamic decompostion of body trajectories. Raw 3D trajectories of two animals can be decomposed into locomotion, non-locomotor movement and body distance. After dynamical temporal decomsition, these three parts are merged together as social behavior motifs for behavioral mapping. **f**, Social behavior metric. Social behavior motifs are clustered and pheonotyped according to the distribution in social behavior space.

After the video acquisition, the multi-animal contour of video capture phase 1 and the single-animal pose of video capture phase 2 are manually annotated for the training of AI to output the 3D poses with identities of animals (Fig. 1b and c). The design of this AI model was separated into four stages for function integration. Through these multistage networks, the task of multi-animal video instance segmentation, pose estimation, and identity recognition was achieved with a relatively small number of manual annotations (∼400 frames), as shown in Fig. 1d (left). By incorporating camera parameters, the above results from various camera angles were combined and matched based on geometric constraints to reconstruct 3D pose trajectories with identities for each animal (Fig. 1d, center and right).

After conducting pose tracking, the process of behavior mapping involves breaking down the trajectories of animals into distinct behavior modules, and then, using appropriate metrics to obtain a low-dimensional representation of these modules^12^. In the context of SBeA, the framework for decomposing behavior is extended from a single animal to multiple animals^12^, with their 3D trajectories being separately decomposed into locomotion, non-locomotor movement, and body distance components (Fig. 1e top and middle). These parallel components are then divided into segments and subsequently merged into social behavioral modules using the dynamic behavior metric (Fig. 1e bottom). Overall, this process utilizes a nature-inspired structure for behavior decomposition and provides a dynamic approach to understanding social behaviors in groups of animals.

To gain insight into the distribution of features within social behavioral modules, it is necessary to convert them into low-dimensional representations (Fig. 1f). These representations incorporate both spatial and temporal aspects, with the spatial aspect being captured by low-dimensional embeddings of distance features in the SBeA framework (Fig. 1f left). The temporal aspect is represented by the social ethogram (Fig. 1f right). In SBeA, social behavioral modules are first clustered based on their spatial characteristics and then expanded into the temporal dimension to construct the social ethogram. This approach allows for a more comprehensive understanding of the distribution of features within social behavioral modules.

### Fewer manual data annotations for multi-animal 3D pose tracking of SBeA

The use of deep learning for social pose estimation has been beneficial in enhancing the acquisition of data for multiple body parts in animals, as previously demonstrated in literature^7,8^. However, due to the flexible social interactions among animals, creating a comprehensive training dataset for deep learning-based social pose estimation is a challenge. Inadequately trained deep neural networks tend to produce higher tracking errors, particularly in frames with close animal interactions ^10^. To address this issue, we introduce a novel animal tracking method using continuously occluded copy-paste data augmentation (Fig. 2a) in our SBeA framework. There are pieces of evidence that simple copy-paste can increase the precision of instance segmentation^15^. Additionally, the continuous copy-paste further increases the performance of multi-object tracking and segmentation^16^. Here for multi-animal tracking, we extend the above methods to continuously occluded copy-paste, which generates the virtual scenario with instance occlusion. By capturing a short video of multiple animals behaving freely in an open field, SBeA obtains sufficient elements (background and animal instances) to generate the virtual dataset. These elements synthesize the complex interactive relationships between animals without the need for manual annotations, resulting in a sufficiently large virtual dataset to train deep neural networks.

**Fig. 2.**
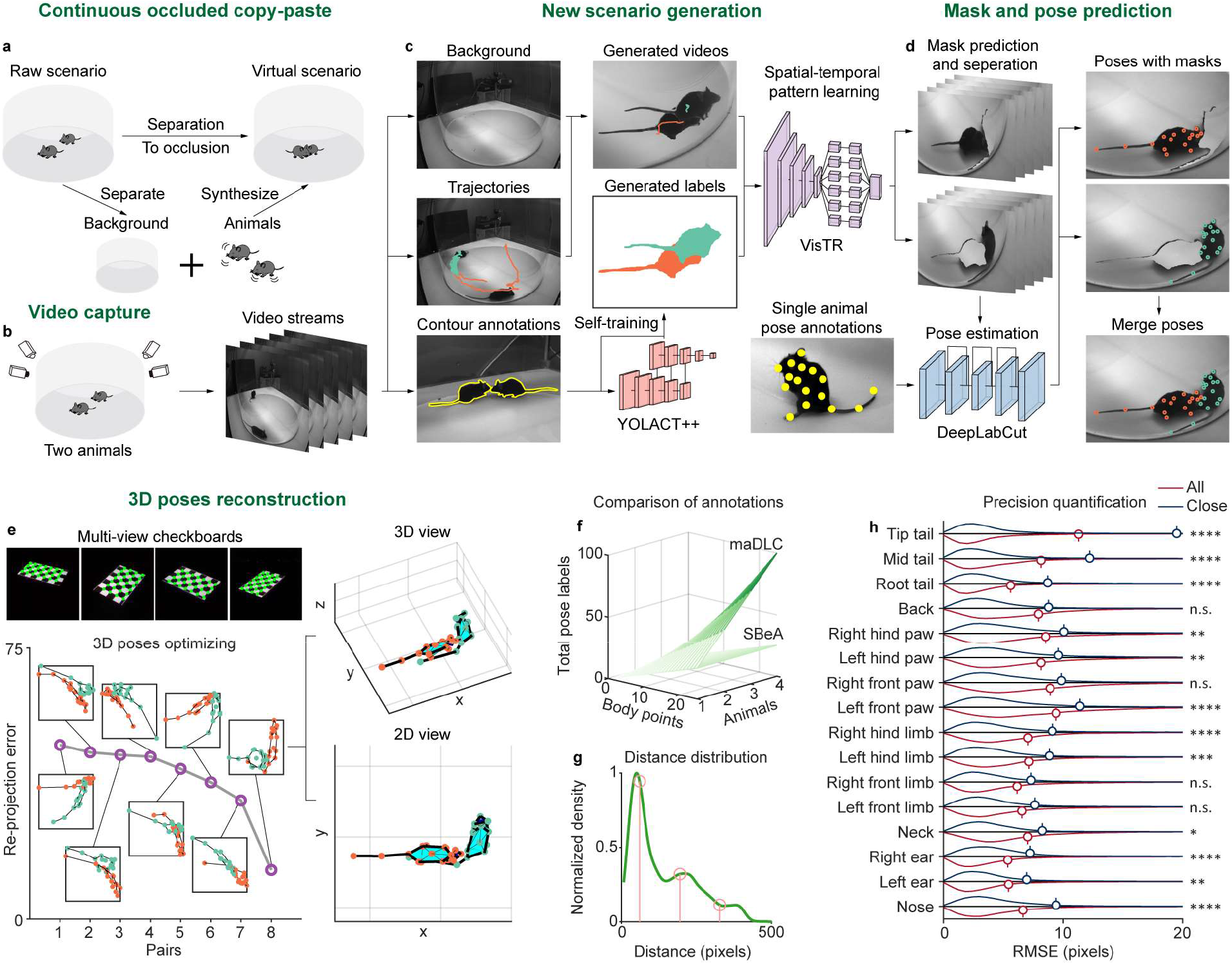
Continuously occluded copy-paste data augmentation-based multi-animal tracking. **a**, Concept diagram of continuously occluded copy-paste data augmentation. From raw scenario, the instances of background and animals can be synthesized with occlusion in new combination. That achieves generating big data from small data. **b**, Video capture of two free-moving animals. Two animals are put in transparent circular open field and the video streams of behavior are captured by camera array. **c**, New scenario generation according to a little manually labeled data. Behavioral video streams are seperated to backgrounds (top left), trajectories (medium left) and manually labeled masks (bottom left). Self-training YOLACT++ is used to predict more unlabeled masks from manually labeled masks. They then combied with backgrounds and trajctories to generate new scenarios of two free-moving mice. **d**, Mask and pose prediction. VisTR is used for the spatial-temporal learning of new scenarios and predict the masks of real mouse instances. Single animal pose estimation model such as DeepLabCut is used for each animal and further the 2D pose of them are merged together. **e**, 3D poses reconstruction. The camera array are calibrated by checkboard images using Zhang’s calibration. And reprojection errors of all combination pairs of 2D poses of each animals are optimized for 3D reconstruction. Top right: 3D view of 3D poses of two mice in this case. Bottom right: 2D view of 3D poses of two mice in this case. **f**, Compasion of the number of manually labeled points of SBeA and maDLC. **g**, Distance distribution of two free-moving mice. Pink stems are distance boundarys clustered by k-means (close: 60.69, interim: 195.03, far: 327.47). **h**, Prediction error compasion of all validataion data. The differences between all and close data are about ±2 pixels (two-way ANOVA followed by Sidak multiple comparisons test). RMSE: root-mean squared error, n.s.: no significant difference, *: P<0.05, **: P<0.01, ***: P<0.001, ****: P<0.0001.

During free behavior, it is common for animals to overlap, leading to loss of tracking in single-view cameras. To address this, SBeA employs a multi-view camera array to capture video streams, enabling compensation for the visual field of the cameras and facilitating continuous tracking (Fig. 2b) ^11–13^. Background and trajectories can be extracted through background subtraction algorithms applied to the raw video streams (Fig. 2c left top and left middle). In addition, frames with close social interactions can be extracted for manual contour annotations (Fig. 2c left bottom). A lightweight instance segmentation deep neural network, YOLACT++, is trained with self-training using approximately 400-800 annotated contour frames (Fig. 2c center bottom), which enhances its performance while ensuring time-efficiency^17,18^. The well-trained self-training YOLACT++ predicts animal masks of video streams, and animal instances can be cropped based on these masks. As some trajectories of two animals may overlap in the same spatial position across different periods, merged animal instances, backgrounds, trajectories, and masks can generate virtual scenarios with various occlusion relationships and mask labels (Fig. 2c center top and center middle). The continuously occluded copy-paste data augmentation increases the scale of the training dataset without additional manual annotations, producing a VIS dataset with successive frames of behaving animals and annotations. To capture the spatial-temporal patterns of occluded animals, the video instance segmentation with transformers (VisTR) method is modified and applied to the virtual VIS dataset as it can segment instances at the sequence level as a whole (Fig. 2c right top)^19^. Well-trained VisTR can patch the raw video streams to display only one animal in each video (Fig. 2d left top and left middle). Thus, pose estimation models trained for single animals, such as DeepLabCut, can be used to predict single animal poses on these patched videos after fine-tuning using patched frames (Fig. 2c right bottom, and 2d left bottom). Finally, the single-animal poses of each patched frame are merged into multi-animal poses (Fig. 2d left top, left middle, and left bottom).

The subsequent step in SBeA, following the acquisition of multi-animal poses from video frames, is the 3D reconstruction (Fig. 2e). Firstly, the MouseVenue3D automatic camera calibration system is employed to acquire the camera parameters of the camera array (Fig. 2e left top)^11,13^. Then, based on the epipolar constraint of camera parameters, the combination of each animal instance in each camera view is optimized to achieve minimum reprojection error (Fig. 2e left bottom). The optimized 3D skeletons of the single frame in Fig. 2d right bottom are presented in Fig. 2e right top and bottom. In the 3D skeleton, the close contact between two animals, such as anogenital sniffing, can be quantified (Fig. 2e right top and bottom).

Compared with the square increasing of routine multi-animal pose estimation methods such as maDLC, the pose annotation strategy in SBeA is linearly increasing with body points and the number of animals (Fig. 2f)^8^. Considering the diversiform social interaction of animals, routine multi-animal pose estimation methods need to annotate more data on frames with various social interactions to get higher precision. Here, we create a well-annotated dataset Social Black Mice for Video Instance Segmentation (SBM-VIS) to quantify the performance of SBeA. According to the distance distribution of the test dataset, the clustering algorithm is used to separate close interaction (Fig. 2g, the left orange stem) and other conditions. The pixel root-mean-square error (RMSE) of all data is significantly lower than the close interaction of about 2 pixels of different body parts (Fig. 2h). But compared with maDLC, SBeA still has significantly lower RMSE of animal close interaction, with 800 pose-labeled frames are used to train maDLC and 400 pose- and 400 mask-labeled frames are used to train SBeA (Extended Data Fig. 1). For all of the test data, SBeA has significantly lower RMSE than maDLC in the Nose, Left ear, Right ear, Root tail, Mid tail, and Tip tail while maDLC has significantly lower pixel RMSE in the Back, Right front paw and Left front paw (Extended Data Fig. 1a). For the close contact part of the test data, SBeA has significantly lower RMSE in Nose, Left ear, Right ear, Left front limb, Right front limb, Left hind limb, Left hind paw, Root tail, Mid tail, and Tip tail (Extended Data Fig. 1b). These results show that SBeA can get higher precision with fewer manual annotations than routine multi-animal pose estimation methods such as maDLC. To get a similar precision of maDLC, SBeA only needs about a quarter of pose annotation points.

### SBeA needs no data annotations for multi-animal identification

Accurately distinguishing the identities of free-moving animals is crucial for social behavior tests, particularly in studying treatment-induced behaviors in transgenic animal models^12,20,21^. However, frequent occlusion of these animals can lead to imprecise identification even with physical markers. Moreover, the animals are the same breed to reduce the influence of irrelevant experiment variables with indistinguishable appearances for human annotators. That causes difficulties in creating the animal identification dataset to train deep neural networks such as SIPEC^22^. To address these challenges, we propose a solution in SBeA, which involves combining a camera array with bidirectional transfer learning in animal identification (Fig. 3a). Transfer learning allows artificial neural networks to use previous knowledge in new tasks^23^. For animal segmentation and identification tasks, the knowledge between them can be shared bidirectionally with each other. So, the segmentation model trained for multi-animals can be transferred to single-animal segmentation, and the identification model trained for single-animals can be transferred to multi-animal identification. The bidirectional transfer learning of them avoids unnecessary manual data annotations.

**Fig. 3.**
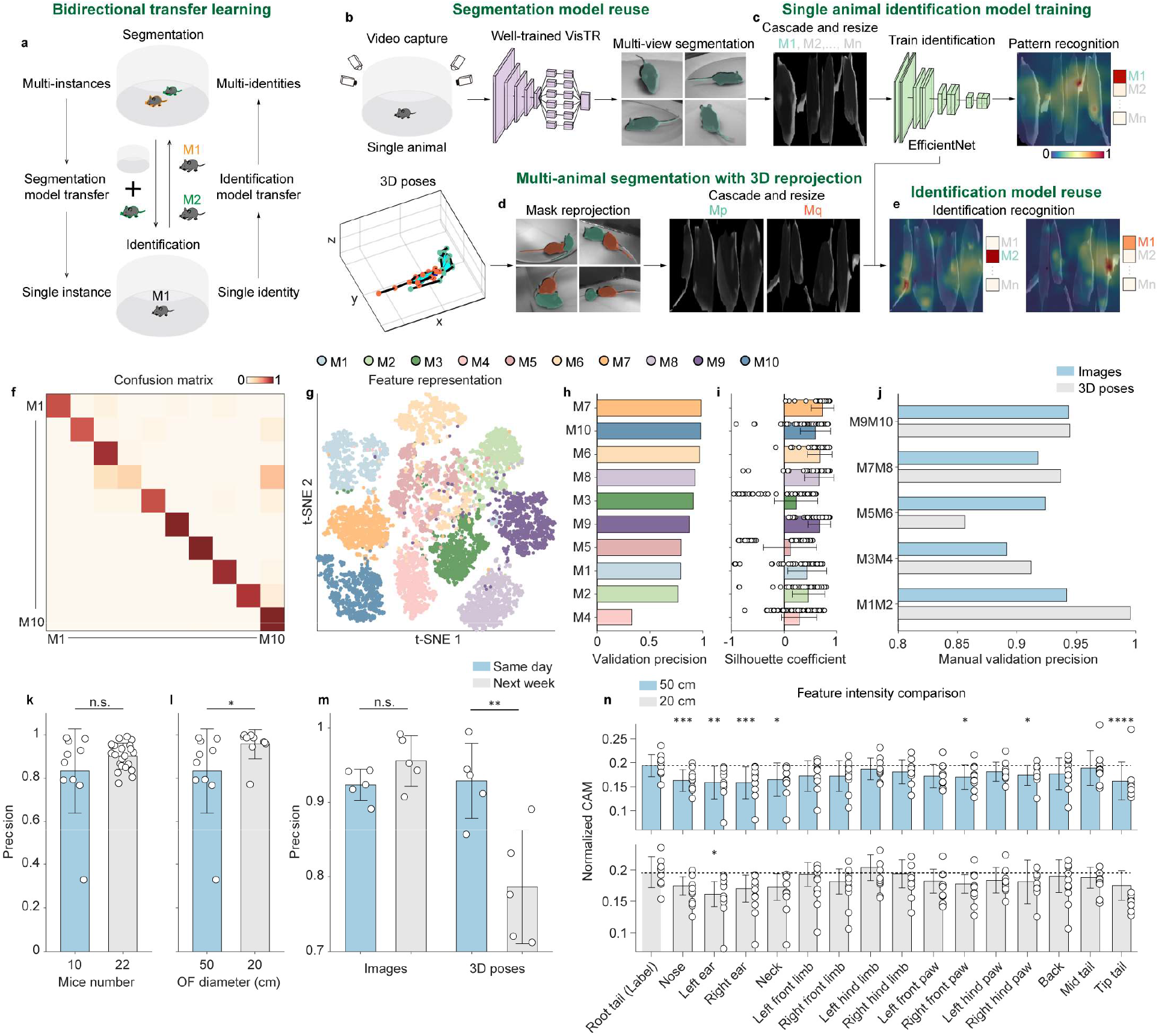
Bidirectional transfer learning-based animal identification. **a**, Concept diagram of bidirectional transfer learning-based animal identification. Well trained segmentation model on multi-animals can be transferred to the single animal, and well trained identity recognition model on the single animal can also be transfered to multi-animals. The transfer learning of two models reduces unnecessary manual annotations of animal identities. **b**, Segmentation model resue. Left: an animal is put in transparent circular open field and the video streams are captured by camera array. Center: The well-trained VisTR is reused for the single animal. Right: The output of well-trained VisTR on the single animal. **c**, Single animal identification model training. Left: the single animal instances of multi-view are cropped, cascaded and resized to an image. Center: using EfficientNet as the backbone to train multi-animal classifiier. Right: The identity recognition pattern visualization by LayerCAM. **d**, Multi-animal segmentation with 3D reprojection. Left: mask reprojection of each camera view. Right: crop, cascade and resize of two animal instances from matched camera view angles. **e**, Identifaction model reuse. The well-trained identifaction model on the single animal can be reused in multi-animal identifaction. **f**, Confusion matrix of single animal identification. **g**, Feature representation of single animal identification using t-SNE. **h**, The sorted validation precision of **f. i**, The sorted silhouette coefficient of **g. j**, The manual validation precision of multi-animal identification. **k**, The identification precision under different mice number. The identification of 10 mice uses 7200 frames for training and 1800 for validation, and 22 mice uses 21600 frames for training and 5400 frames for validation. With the increase of animal number, the add of training frames can keep higher identification precision (two-sided Mann–Whitney test). **l**, The identification precision under different open field (OF) diameter of behavioral test(two-sided Mann–Whitney test). **m**, The identification precision in different interval time between social behavior test and identify recording(two-sided unpaired T-test). **n**, The feature intensity of the tracking body parts under different OF diameter of behavioral test. The root tail of mice is labeled by different black line markers for the easy distinguish of human(one-way ANOVA followed by Dunnett multiple comparisons test). n.s.: no significant difference, *: P<0.05, **: P<0.01, ***: P<0.001, ****: P<0.0001.

Well-trained VisTR in Fig. 2 can be used to segment single-animal instances from multiple view angles (Fig. 3b). These instances are then cropped, cascaded, and resized to generate training data for an identification model based on EfficientNet architecture (Fig. 3c, left and center)^24^. After that, LayerCAM is used to evaluate the patterns for identification recognition (Fig. 3c right)^25^. Before using the identification model in multi-animal instances, the cascaded and resized image frames were prepared (Fig. 3d, right). By using the best geometric constraint of 3D poses, instances from each frame view angle of each animal were matched to construct input frames of the identification model (Fig. 3d, left). Finally, the well-trained model outputted the top prediction probabilities to append the identities of instances and 3D poses. LayerCAM was also employed to verify the recognition patterns for identification (Fig. 3e).

To evaluate the identification model’s performance in SBeA, we conducted experiments with ten C57BL/6J mice having tail markers, where we recorded their free behaviors for 5 minutes. The tail markers are convenient for experimenters to distinguish the identities of each mouse. The first 4 minutes of data were used for training the identification model, and the last 1 minute was used for validation. The confusion matrix of the validation data demonstrated that the EfficientNet model can identify most of the mice (Fig. 3f). The t-SNE algorithm was used to create a 2D feature representation of the identified mice (Fig. 3j). However, the features of mice with ID M4 and M5 were found to be mixed with other classes, as quantified by the silhouette coefficient Fig. 3h). The sorted validation precision of the identified mice showed that the mouse with ID M4 had the lowest precision of approximately 0.4 (Fig. 3i). Even though the features of M5 were mixed with other classes, its precision was found to be around 0.8 (Fig. 3i).

To assess the identification model’s performance in multi-animal data, we recorded the free social behaviors of 5 paired C57BL/6J mice identified by SBeA for 15 minutes. We manually verified the identities of mask reprojection images and 3D poses frame by frame (Fig. 3j). The results indicated that although some of the single mouse identity precisions were lower (Fig. 3i), the overall precision in identifying pairs of mice could be higher than 0.85, as seen in the case of the pairs of M3&M4 and M5&M6. Additionally, the validation precision in single-animal identification was found to be positively correlated with precision in multi-animal identification, as evidenced by the other pairs (Fig. 3j).

We also investigate if the number of animals would influence the identification recognition precision. Previous research suggests that the identification precision may decrease with an increasing number of animals involved in the study^26,27^. To counteract this trend, we increased the amount of training data to balance the precision decrease. Our results indicate that for a group of 22 mice, a 15-minute video recording can achieve similar precision to that of 10 mice with a 5-minute recording (Fig. 3k). These findings have important implications for optimizing study design and ensuring accurate identification of individual animals in social behavior experiments.

Our research has revealed that the precision of animal identification can be influenced by the experiment apparatus used in social behavior tests (Fig. 3l). Specifically, we found that open fields with different diameters -50cm and -20cm can impact the precision of animal identification conducted on the same ten mice. Our results indicate that the precision of identification in the 20cm open field is significantly higher than that in the 50cm field (Fig. 3l). This difference may be due to the higher dots per inch (DPI) of mice.

Further, we tested the stability of identification patterns. Animals would groom themselves, which could change the patterns of identities^9^. We compared the identification precision of two separate groups of mice. One group underwent both identity video recording and social behavior tests on the same day, while the other group underwent social behavior tests one week after their identity videos were recorded (Fig. 3m). We manually verified the identities of mask reprojection images and 3D poses frame-by-frame. Our analysis revealed that while there was no significant difference in the precision of mask reprojection images between the two groups, the precision of 3D poses in the group that underwent social behavior tests one week after the recording of their identity videos was significantly lower than that of the group that underwent both on the same day (Fig. 3m). As the precision of 3D poses is equivalent to the identification precision of cascade and resize images, the observed decrease in precision of 3D poses indicates a decline in identification precision. Shorter intervals between the recording of identity videos and social behavior tests could potentially enhance the accuracy of identification recognition.

We evaluated the feature intensity of the identification model used to distinguish different animals at last in this chapter. To this end, we designed open fields with diameters of 50 and 20 cm, respectively, in which the same ten mice were allowed to freely engage in social behavior 2 mice per trial. The pose tracking point “Root tail” with tail markers was used as a control against other body parts (Fig. 3n). We calculated LayerCAM values to quantify the feature intensity of each body point. The results showed that the Root tail in the 50 cm group had more significant feature intensities than in the 20 cm group. This finding suggests that a higher DPI can enable the identification model to capture more available fur pattern features and thereby overcome errors resulting from marker occlusion. Additionally, we found that identification using low animal DPI requires the use of stronger markers to maintain sufficient recognition precision.

### SBeA reveals the social behavioral structure in the atlas by unsupervised machine learning

Following pose tracking, it is necessary to map the trajectories with animal identities to a low-dimensional space to gain insights into behavior (Fig. 4a). Recent research has indicated that the body language of social animals can be represented through sequential behavioral motifs or modules^28^. Thus, we expand our prior work on the animal behavior mapping framework to encompass multiple animals, Behavior Atlas (BeA), which was initially developed for a single animal. The concepts of parallel and dynamic behavior decomposition from BeA have been adopted in our new framework SBeA (Fig. 4b and c). In the social process, the distance between animals is an essential component, as noted in previous studies^29^. In addition to using non-locomotor movement to assess body movement and locomotion to evaluate body displacement, body distance is also utilized to evaluate the relationships of body position (Fig. 4b). After parallel decomposition, each component is decomposed further using dynamic time alignment kernel (DTAK) to retain the natural dynamic structures of behavior (Fig. 4c). To distinguish subtle structures of social behavior, the temporal points of decomposition for each component are merged through logical addition (Fig. 4d). The aforementioned steps enable the metric of social behavior, resulting in the transformation of continuous pose trajectories into discrete social behavior modules.

**Fig. 4.**
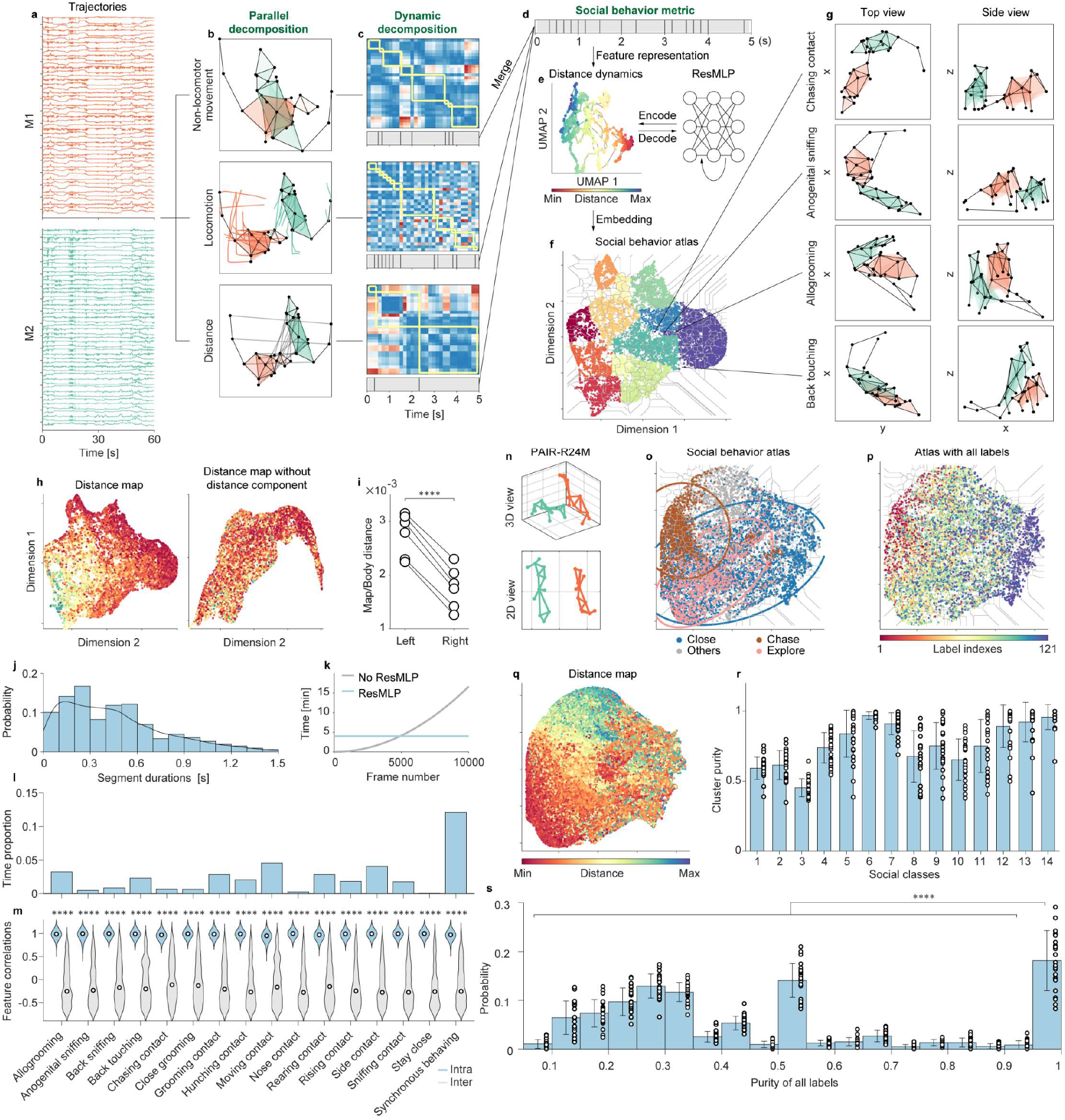
Natural behavioral structure-inspired segmentation and mapping of free social interactions. **a**, The 3D trajectories of 2 animals. **b**, The parallel decomposition of trajectories. Top: Non-locomotor movement. Middle: Locomotion. Bottom: Distance. **c**, The dynamic decomposition after parallel decomposition using Dynamic Time Alignment Kernel (DTAK). **d, e**, and **f**, Social behavior metric after dynamic decomposition. **d**, Decomposed segments merging. **e**, Feture representation of segments. Left: Dimensional reduction of distance dynamics. Right: Residual Multi-Layer Perceptron (ResMLP) for feature refining. **f**, Social behavior atlas construction. The adaptive watershed is used for clustering. Color represents large clusters and area represent sub-clusters. **g**, Social behavior cases clustered in social behavior atlas. **h**, Left: The distance map of **f**, Right: The distance map without the distance component in behavior decomposition steps. **i**, The comparison of map/body distance of **h** (two-sided paired T-test). The higher the map/body distance, the better performance for the representation of social behavior with different distances. **j**, The probability of segment durations. **k**, The comparison of computational time consumption of feature representation with or without ResMLP. **l**, The time proportion of different behavior. **m**, The feature correlations intra and inter behavioral classes (two-sided Mann–Whitney test). **n-s**,The performance quantification of SBeA on the PAIR-R24M dataset. **n**, The visualization of two mice in the PAIR-R24M dataset. **o**, The social behavior atlas of PAIR-R24M dataset. The social classes of the PAIR-R24M dataset are seperated in social behavior atlas. The ellipse is the Gaussian model fitting of the three classes. **p**, The social behavior atlas of all the class labels of PAIR-R24M dataset. The 11 classes of each mouse are combined to 121 classes, and the 121 classes are distributed with patterns. **q**, The distance map of social behavior atlas. The distance distribution of distance map is coincident with labels in **o. r**, The cluster purity of social classes in **o. s**, The cluster purity probability of all labels in **p**. The cluster purities greater than 0.95 are significant higher than others (one-way ANOVA followed by Tukey multiple comparisons test). n.s.: no significant difference, *: P<0.05, **: P<0.01, ***: P<0.001, ****: P<0.0001.

Then, the social behavior modules are embedded in a low-dimensional space for behavior representation (Fig. 4e and f). All of the social behavior modules from different experimental trials need to be represented in a common feature space. That induces two questions, the first is what feature is reasonable to represent social behavior in a low-dimensional space, and the second is how to create a common feature space under the big behavioral data^30,31^. For the first question, the distance component is chosen for the feature representation of social behavior modules (Fig. 4e left). The dimensionally reduced distance component by uniform manifold approximation and projection (UMAP) is beneficial to improve the separation of behavior atlas verified by our previous studies^11–14,32^. But with the increase of data scale, the computational consumption of UMAP would be unacceptable because of limited memory space, which is the second question. To solve the second question, the residual multilayer perceptron (ResMLP) is combined with UMAP for feature representation (Fig. 4e right)^33^. A part of the social behavior feature frames is extracted randomly to build up the feature representation of distance dynamics by the UMAP. Then, the mapping from extracted social behavior feature frames to distance dynamics is trained by ResMLP for the feature encoding. Further, the rest of the social behavior feature frames are decoded by ResMLP to distance dynamics. The distance dynamics are embedded by DTAK and UMAP to construct the social behavior atlas (Fig. 4f). To reveal the distributions of different social behavior modules, based on density clustering, we modified the watershed algorithm to automatically determine the best cluster density with upper and lower boundaries. At last, the social behavior modules of the same clusters are manually identified and defined (Fig. 4g).

In constructing the social behavior atlas, the inclusion of the distance component is crucial. By using the distance component, the social behavior atlas can maintain the overall body distance structures of social behavior modules (Fig. 4h left), while the absence of the distance component leads to a lack of observable patterns in the distribution of distance (Fig. 4h right). To compare the effectiveness of the distance representations in the atlases, the map/body distance metric is utilized, with higher values indicating better performance in distance representation (Fig. 4i). Results show that the distance component is essential in achieving a high map/body distance, indicating the importance of including this component in constructing the social behavior atlas. Additionally, the 0.45±0.32s temporal duration of merged behavioral modules reveals that the SBeA framework can effectively decompose social behavior into dynamic sub-second motifs (Fig. 4j)^12,34^. The ResMLP can address issues related to the memory cost of large behavioral data, while also reducing computational time consumption compared to using UMAP alone. More than 5000 frames can get time benefits from ResMLP, and the time benefits will increase with the number of frames (Fig. 4k). Then, the time proportion of identified behavioral modules is quantified to evaluate their temporal precision (Fig. 4l). The time proportion of the typical social behavior such as allogrooming conforms to previous studies on social behavior^35^. Further, the feature correlations between the intra- and inter-clusters of each social behavior class are compared for the evaluation of clustering consistency (Fig. 4m). The intra-feature correlations of each social behavior class are significantly higher than inter-feature correlations, and the intra-feature correlations distribute consistently near to 1, in turn, the inter-feature correlations distribute in the weak negative correlation. These unsupervised validation measures demonstrate the effectiveness of the SBeA framework in accurately mapping social behavior.

In addition to unsupervised validation, we conducted supervised validation of SBeA using the PAIR-R24M dataset (Fig. 4n)^36^. The dataset provides 3D poses, social behavior labels, and subject behavior labels of rats in free behavior. We used SBeA to construct the social behavior atlas for the dataset, and appended the three social labels (close, chase, and explore) to each behavior module (Fig. 4o). The distributions of the three social labels were separated and matched their similarity relationship. The 121 combinations of subject behavior labels also showed distribution patterns in the social behavior atlas (Fig. 4p). The social labels such as close and explore were consistent with the close distance distribution in the distance map, and the chase label was consistent with the distance transition zone of the distance map (Fig. 4q). To quantify the clustering performance, we used the cluster purity of social and subject behavior labels (Fig. 4r and s). For the upper boundary of clustering, 14 classes were clustered with a mean cluster purity of 0.77±0.16 (Fig. 4r). For the lower boundary of clustering, 405 classes were clustered, and the probability of cluster purities greater than 0.95 was significantly higher than for other purities (Fig. 4s). These results provide further validation of the performance of SBeA in supervised contexts.

### SBeA identifies *Shank3B* knockout mice in free-social interactions by subtle behavior modules

Social behavior can serve as an indicator of the genetic variations that underlie neuropsychiatric disorders^37^. SBeA is well-suited for this purpose, as it allows for a detailed characterization of social behavior at an atlas-level. To test the ability of SBeA to detect genetic differences from social behavior, we utilized an animal model of autism spectrum disorder (ASD): Shank3B knockout mice^12,20^. While abnormal individual behaviors of these mice have been previously identified, the limitations of existing techniques have made it difficult to fully understand their abnormal free social behaviors^12,20^.

To distinguish between Shank3B knockout (KO) mice and wild-type (WT) mice, a free-social behavioral paradigm was designed based on the framework of SBeA, which consists of three steps: identity recording, social behavior recording, and SBeA processing (Fig. 5a). First, the home-caged WT and KO mice were randomly shuffled and recorded for 5 minutes each using the MouseVenue3D system to obtain identity information. After identity recording, the mice were randomly grouped into three pairs (WT-WT, WT-KO, and KO-KO) for social behavior recording, with each pair of mice recorded for 15 minutes. The identity and social behavior data were then processed using SBeA for 3D pose tracking and behavior mapping. The experiment used a total of 10 WT and 10 KO mice, resulting in 45 unique pairs of mice, including 10 WT-WT, 10 KO-KO, and 25 WT-KO pairs. To ensure equal representation of each group, the number of WT-KO pairs was reduced from 25 to 10 through random sampling. Before behavior mapping, the raw trajectories were copied and switched to capture the direction of social behavior between WT and KO mice. This resulted in a total of 60 pairs of trajectories for behavior mapping using SBeA.

**Fig. 5.**
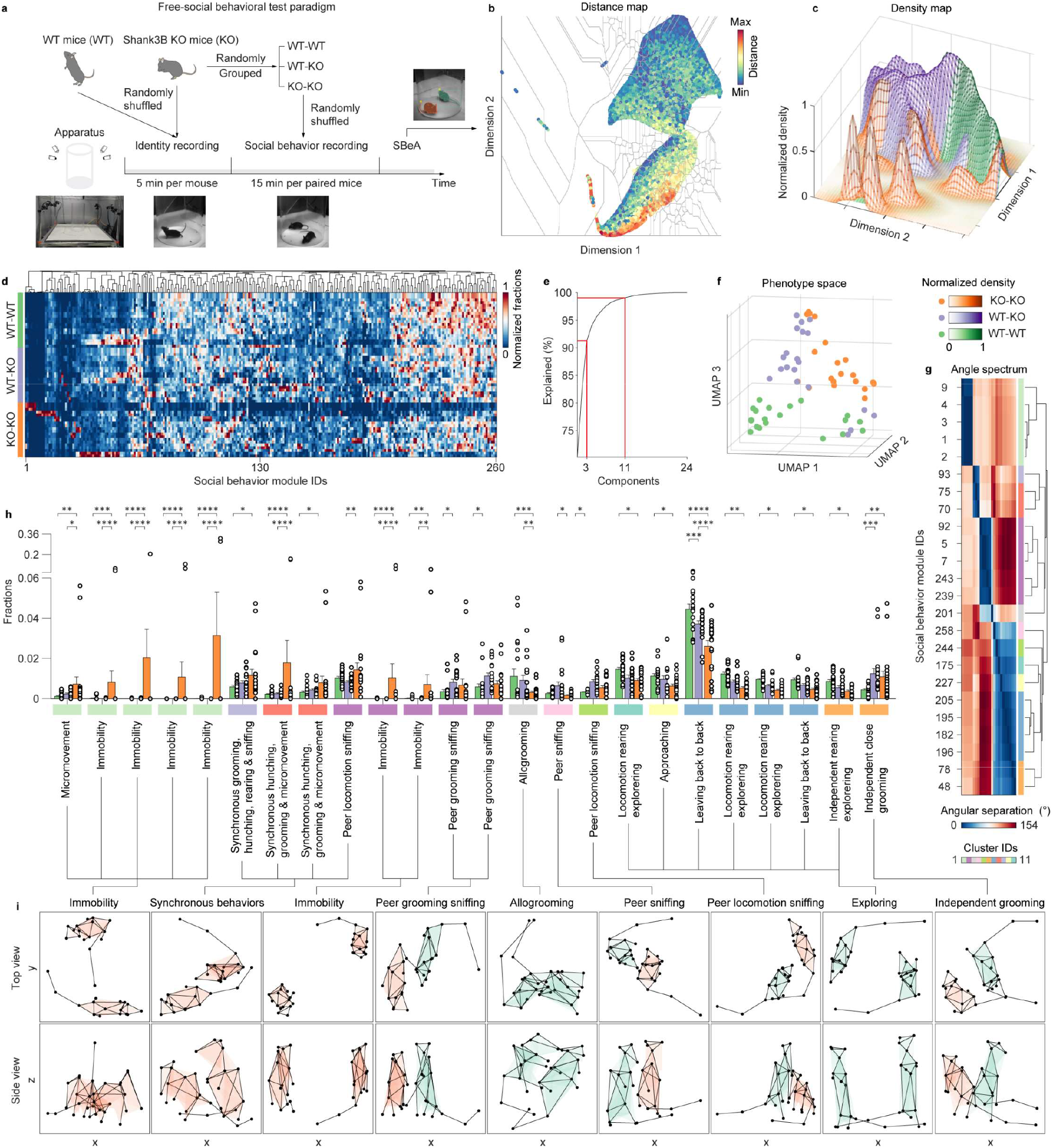
The identifying of abnormal social behavior modules in *Shank3B* knockout mice. **a**, The paradigm of free-social behavioral test. WT: Wild type mice, KO: *Shank3B* knockout mice. **b**, The social behavior atlas with distance map of 3 grouped mice. **c**, The distribution of social behavioral modules of three social groups. A total of 260 social behavior modules are identified. **d**, The fractions of social behavioral modules of three social groups. The fractions of each group are normalized, and they are clustered and resorted according to the dimension of social behavior modules. **e**, Dimensional reduction of behavior fractions using principal component analysis (PCA) after hypothesis testing (two-way ANOVA followed by Tukey multiple comparisons test). 24 social behavior modules are significant differences in three groups. 3 components can explain more than 90% variances, and 11 components can explain more than 99% variances. **f**, The construction of phenotype space. UMAP is used to reduce the 260 dimensions of social behavior modules to 3 dimensions according to **e**. Different colors of dots represent different social groups. The phenotypes of 3 social groups can be seperated in phenotpye space. **g**, The merging of social behavior modules according to behavioral feature angles and **e**. 24 social behavior modules with significant differences are firstly mapped to PCA feature space and then the angular separation are calculated to construct angle spectrum. Further, hierarchical clustering is used to cluster angle spectrum to 11 clusters according to **e. h**, The comparison of beahvioral fractions of 3 social groups. 24 social behavior modules with significant differences are manually identified. **i**, The visualization of merged social behavior modules. With the assistance of **g**, 9 social behavior modules are merged and identified from 24 social behavior modules. Orange 3D mice represent KO mice, and green 3D mice represent WT mice. n.s.: no significant difference, *: P<0.05, **: P<0.01, ***: P<0.001, ****: P<0.0001.

The social behavior atlas with distance map is shown in Fig. 5b. After the construction of the social behavior atlas, the density map is calculated to compare the social behavior distribution of each group by kernel density estimation (Fig. 5c). Density map shows obvious differences across the three groups. Combing with the distance map, the WT-WT group shows social behavior phenotypes with flexible distances from close to far, the KO-KO group shows more abnormal social behaviors than the WT-WT group, and WT-KO shows more close social interaction than the WT-WT group. From the global level, the social behaviors of KO mice show differences from WT mice.

The 260 social behavior modules identified in the social behavior atlas were clustered to reveal their coincident patterns, which displayed distinct speckled patterns for each group, ranging from 1 to 20 social behavior modules in the KO-KO group (Fig. 5d). To compare the differences in behavior components among the three groups, principal component analysis (PCA) was employed to determine the percent variability explained by each principal component (Fig. 5e). The results indicated that three components could account for 90% of the variance, while 11 components could account for 99% of the variance. To construct the phenotype space of the three groups, UMAP was used for dimensional reduction of the social behavior modules, with the dimension number set to 3 based on the 90% variance explanation, owing to the more robust feature representation of non-linear dimensional reduction (Fig. 5f). The distributions of the three groups in the phenotype space were found to be segregated, matching the distribution of the density map, and distinguishing KO mice from WT mice (Fig. 5c).

Further, SBeA was utilized to identify subtle social behavior modules that distinguish KO and WT mice. The two-way ANOVA was used to compare the behavior fractions between the three groups, and 24 social behavior modules were found to have significant differences (Fig. 5h). To reduce the redundancy of these results, angle spectrum clustering, which combines PCA and hierarchical clustering, was proposed (Fig. 5g). The social behavior modules were merged based on their angular separation of features, resulting in the identification of 9 social behaviors, as determined by human analysis (Fig. 5i). The color of mice represented the behavior cases with the highest mean fraction in Fig. 5g.

The 9 social behavior modules identified through SBeA highlighted significant differences among the three groups. The WT-WT group exhibited more allogrooming, a prosocial behavior, than the WT-KO and KO-KO groups^38^. Conversely, allogrooming was rare in unstressed partners and even rarer in Shank3B KO mice, suggesting an antisocial behavioral phenotype^35^. The exploring behavior of the WT-WT group was significantly higher than that of the KO-KO group, which displayed reduced motor ability or social novelty^12,20^. In the WT-KO group, social behavior with significant differences were divided into two parts, namely, peer sniffing and independent grooming. Peer sniffing was observed more frequently in the WT mouse, particularly when the KO mouse was grooming or in locomotion, indicating a behavioral phenotype of curiosity. Furthermore, the KO mouse could induce higher interest in the WT mouse than vice versa. Independent grooming could be an imitation of the WT mouse by the KO mouse, and in the KO-KO groups, the higher incidence of independent grooming could be attributed to the increased individual grooming of each mouse. In addition to increased independent grooming, two abnormal behavior phenotypes, namely, synchronous behaviors and immobility, were observed. The synchronous behaviors displayed 5 subtypes, including grooming, hunching, rearing, sniffing, and micromovement, indicating greater behavior variability in free-social conditions compared to individual spontaneous behavior of KO mice^12^. Most instances of immobility occurred in only one pair of KO-KO mice, indicating that abnormal autistic-like behaviors vary even among mice with the same genetic background. These findings demonstrate that SBeA can differentiate genetic mutant animals based on social behavior and identify genetic mutant-related subtle social behavior modules.

### SBeA is robust to be used in different environments across species

To assess the generalizability of SBeA to different animal species and experimental settings, the behaviors of birds and dogs were captured using the MouseVenue3D system with varying device configurations^11^. The animals were prepared to have as similar appearances as possible (Fig. 5a top and 5e top), and it was difficult for human experimenters to separate two animals from the randomly selected frames. The resulting videos were manually annotated to train the AI of the pose tracking component of SBeA (Fig. 6a bottom and 6e bottom), using 19 body parts for birds and 17 body parts for dogs, based on previous studies^39,40^. The well-trained AI was then used to predict video instances, body poses, and identities (Fig. 6b and f), which were mapped to a social ethogram and behavior atlas using the behavior mapping component of SBeA (Fig. 6c and g). In total, 34 and 15 social behavior classes were identified for birds and dogs, respectively, and their typical cases were visualized in 3D (Fig. 6d and h). The 3D pose tracking of birds showed clear identification of their claw touching their rectrix, while the 3D pose tracking of dogs was robust to occlusion even in the lying posture.

**Fig. 6.**
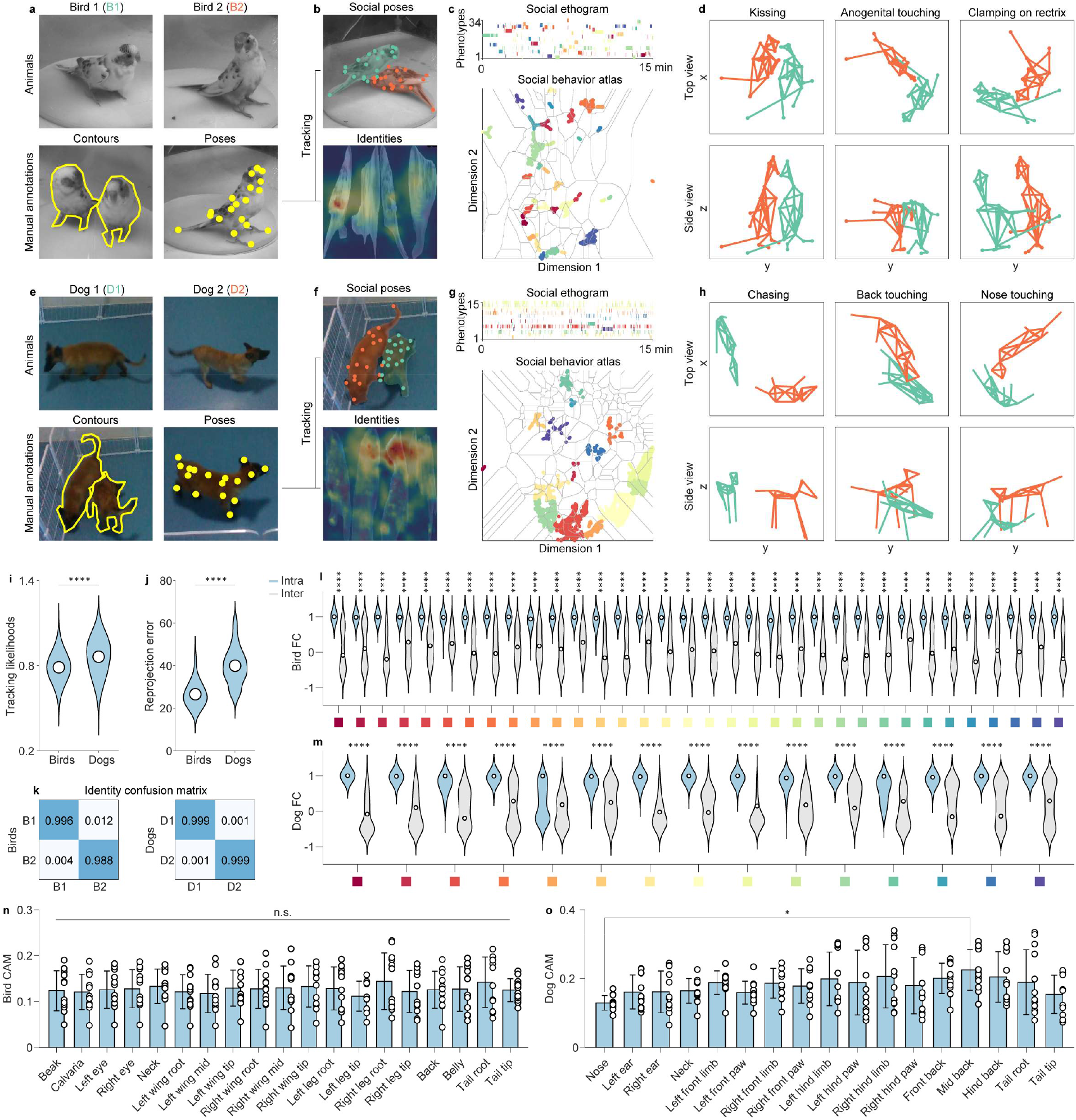
SBeA for the applications across species such as birds and dogs. **a-d**, SBeA is used for birds. **a**, The preparation of birds. Two parrots with inconspicuous appearance difference are used for social behavior test. After video recording of identiy and free-social behavior by camera array, the contours and poses are manually annotated. 19 body parts are defined for 3D pose tracking. **b**, The social poses and identities outputs of SBeA. **c**, The social ethogram and social behavior atlas of birds. **d**, The 3D social behavior cases of birds. **e-h**, SBeA is used for dogs. **e**, The preparation of dogs. Two Belgian Malinois with inconspicuous appearance difference are used for the social behavior test. After video recording of identity and free-social behavior by camera array, the contours and poses are manually annotated. 17 body parts are defined for 3D pose tracking.. **f**, The social poses and identities outputs of SBeA. **g**, The social ethogram and social behavior atlas of dogs. **h**, The 3D social behavior cases of dogs. **i-o**, The performance quantification of SBeA in birds and dogs. **i**, The tracking likelihoods of birds and dogs are significant different (two-sided Mann–Whitney test). **j**, The 3D reprojection error of birds and dogs are significant different (two-sided Mann–Whitney test). **k**, The identity recognition confusion matrix of birds and dogs. **l**, The feature correlations (FC) intra and inter behavioral classes of birds (two-way ANOVA followed by Sidak multiple comparisons test). **m**, The FC intra and inter behavioral classes of dogs (two-way ANOVA followed by Sidak multiple comparisons test). **n**, The feature intensity of the tracking body parts of birds (one-way ANOVA followed by Tukey multiple comparisons test).The feature intensities do not show significant differences. **o**, The feature intensity of the tracking body parts of dogs (one-way ANOVA followed by Dunnett multiple comparisons test).The feature intensities between nose and mid back show significant differences. n.s.: no significant difference, *: P<0.05, **: P<0.01, ***: P<0.001, ****: P<0.0001.

To evaluate the performance of the SBeA algorithm in tracking birds and dogs, various metrics were employed, including tracking likelihood, 3D reprojection error, identity confusion matrix, and feature correlation (FC) (Fig. 6i-m). The results indicate that while dogs have a higher tracking likelihood than birds, both achieve a satisfactory level of tracking precision (Fig. 6i)^12^. But the 3D reprojection error is significantly higher for dogs due to incomplete camera coverage and annotation errors (Fig. 6j). In terms of identity recognition, both birds and dogs have higher precision than mice due to their distinct fur patterns (Fig. 6k). The results of FC show that all of the intra-FC of clusters are significantly higher than inter-FC (Fig. 6l and m). But from the distribution of FCs, the clustering performance of birds is better than dogs. The feature mix-up of intra- and inter-clusters is influenced by the 3D pose tracking precision. The error of 3D pose tracking such as target loss in dogs would degrade the performance of SBeA clustering. The LayerCAM analysis reveals no significant differences in feature values between birds and dogs, except for the Mid back and Nose of dogs, which may be attributed to the loss of nose detection in video captures (Fig. 6n and o). The identification recognition network automatically reduces the feature weights on the body part with target loss or occlusion to keep the higher recognition precision of identities. These results demonstrate that SBeA is robust enough to be applied to different animal species in various experimental settings, making it a versatile tool for the study of social behavior in animals.

## Discussion

Here we have presented SBeA, a framework for 3D pose tracking and behavior mapping of multiple free-social animals. SBeA builds upon the BeA framework, extending it to enable multi-animal pose estimation and social behavior clustering^11–14^. The method reduces the labor required for annotation by up to fifty percent compared to traditional approaches for pose estimation. By utilizing four cameras, SBeA overcomes the issue of occlusion and reconstructs 3D behaviors accurately. Notably, SBeA resolves the challenge of animal identification over extended time frames, facilitating the study of close social interactions. The technique is highly versatile and has been successfully applied to various animal species, including *Shank3B* knockout mice, where it revealed synchronous behaviors and reduced social interest. SBeA’s cross-species application has been verified in birds and dogs. In summary, SBeA represents a breakthrough in deep learning-based pose estimation and identification, offering numerous potential applications in animal behavior research.

Both maDLC and SLEAP are versatile tools that can be applied to a variety of animal models, from fish to humans.^7,8^. However, a major drawback of these tools is the lack of a framework for maintaining animal identities during long-term experiments, which can be fatal to the accuracy of results^10^. SBeA incorporates the identity recognition approach of idTracker.ai and TRex, utilizing deep neural networks to directly learn the appearance features of animals^26,41^. This results in a lower error rate than maDLC or SLEAP and allows for frames with low accuracy to be filtered without affecting the entire video. Additionally, SBeA provides an extension of 2D tracking tools to 3D movement tracking, which is critical for making accurate inferences about animal behavior.

One potential area for future research to improve SBeA is the development of an end-to-end model that can reduce storage consumption. To accomplish this, the process of data generation could be incorporated into a video instance segmentation model. Additionally, the identity videos available in this context may contain sufficient information to train a deep learning model for tasks such as multi-animal segmentation, identification, and pose estimation. Furthermore, the behavior atlas of a single animal could be combined with a social behavior atlas of multiple animals through an algorithmic bridge from BeA to SBeA that facilitates not only social behavior analysis but also other forms of analysis within the field.

## Online content

The online version of SBeA will be released after the peer review of this work. Anyone interested in our work can contact us for the further corporation.

## Methods

### Experiments of mice, birds, and dogs

There are four experiments in this study.

The first is the free-social behavior test of two wild-type mice for the program design of SBeA. 32 adult male C57BL/6 mice (7–12 weeks old) are used for the free-social behavior test. The mice were housed at 4-5 mice per cage under a 12-h light–dark cycle at 22–25 °C with 40–70% humidity, and were allowed to access water and food ad libitum (Shenzhen Institutes of Advanced Technology, Shenzhen, China). Before the social behavior test, the mice are added tail tags using black mark pen. The tail tags are constructed by horizontal and vertical lines. The horizontal line represents one, and the vertical line represents five. Using the combination of horizontal and vertical lines, the mice are marked according to the sequence of the experiment. After that, the mice are put into a circular open field made of a transparent acrylic wall and white plastic ground, with a base diameter of 50 cm or 20 cm and a height of 50 cm for 5 min or 15 min identity recording one by one using MouseVenue3D. Then, the mice are paired and put into the same circular open field for the free-social behavior test.

The second is the free-social behavior test of mice with different genotypes. 5 adult (8 weeks old) Shank3B knockout (KO; *Shank3B*^−*/*−^) mice on C57BL/6J genetic background and 5 adults (8 weeks old) male C57BL/6 mice, were used in the behavioral experiments. *Shank3B*^−*/*−^ mice were obtained from the Jackson Laboratory (Jax No. 017688) and were described previously^20^. The mice were housed at 4-5 mice per cage under a 12-h light–dark cycle at 22–25 °C with 40–70% humidity, and were allowed to access water and food ad libitum (Shenzhen Institutes of Advanced Technology, Shenzhen, China). The mice have added the tail tag introduced above. After that, the mice are put into a circular open field with a base diameter of 20 cm introduced before for 5 min identity recording. Then the mice are paired to WT-WT, WT-KO, and KO-KO groups and put into the same circular open field for the free-social behavior test. The combinations of groups and the sequence of experiments are random generated by customized MATLAB code.

The third is the free-social behavior test of two birds. One male and one female Melopsittacus Undulatus (about 26 weeks old) are used in this experiment. They are housed in a conventional environment with feed regularly (Shenzhen Institutes of Advanced Technology, Shenzhen, China). The birds are first put into a circular open field with a base diameter of 20 cm introduced before for 5 min identity recording one by one, and then put in it together for 15 min free-social behavior test and recording.

The fourth is the free-social behavior test of two dogs. Two female Belgian Malinois (13 weeks old) are used in this experiment. They are housed in Kunming Police Dog Base of the Chinese Ministry of Public Security, Kunming,650204, China, and their behavior test of them is finished in the State Key Laboratory of Genetic Resources and Evolution, Kunming Institute of Zoology, Chinese Academy of Sciences, Kunming, 650223, China. The dogs are first put into a 2 × 2 m^2^ open field made by fences one by one for the identity recording. Restricted by the locomotion of dogs, there are only 6 min and 11 min identity frames captured by MouseVenue3D and both of them are used for identification. Then, they are both put into the open field for 15 min free-social behavior test.

All husbandry and experimental procedures of mice and birds in this study were approved by Animal Care and Use Committees at the Shenzhen Institute of Advanced Technology, Chinese Academy of Sciences. And all husbandry and experimental procedures of dogs in this study were approved by Animal Care and Use Committees at the Kunming Institute of Zoology, Chinese Academy of Sciences.

### MouseVenue3D subtle behavior capture system

There are three versions of MouseVenue3D systems used in this study.

The first version is used for the data capture of the SBM-VIS dataset. Four Intel RealSense D435 cameras are mounted orthogonally on four supporting pillars made of stainless steel. The distance between the nearest cameras is 90 cm. The cameras are adjusted to 75 cm off the ground to capture the whole view of the animal activities in the open field. Images were simultaneously recorded at 30 frames in 640×480 sizes per second by a PCI-E USB-3.0 data acquisition card and the pyrealsense2 Python camera interface package. The cameras are connected to a high-performance computer (i7-9700K, 16G RAM) equipped with a 1-terabyte SSD and 12-terabyte HDD as an image acquisition platform. The computer also controls the camera calibration module.

The second version is used for the behavioral capturing of mice and birds. Four Point Grey FLIR Chameleon3 CM3-U3-13S2 cameras with adaptive zoom lenses are mounted orthogonally on four supporting pillars made of stainless steel. The distance between the nearest cameras is 85 cm. The cameras are adjusted to 45 cm off the ground to capture the whole view of the animal activities in the open field. To adapt to the size of the open field, the focal length and the pitch angle of cameras are flexibly adjusted before each experiment. Images were simultaneously recorded at 30 frames in 1288 ×964 sizes in grayscale per second by a PCI-E USB-3.0 data acquisition card and the Spinnaker Python camera interface package. The cameras are connected to a high-performance computer (i9-10900K, 128G RAM) equipped with a 512-gigabyte SSD and two 16-terabyte HDDs as an image acquisition platform. The computer also controls the camera calibration module.

The third version is used for the behavioral capturing of dogs. Four Intel RealSense D435 cameras are mounted orthogonally on walls. The distance between the nearest cameras is 210 cm. The cameras are adjusted to 150 cm off the ground to capture the whole view of the dog activities in the open field. Images were simultaneously recorded at 30 frames in 640×360 sizes per second by a PCI-E USB-3.0 data acquisition card and the pyrealsense2 Python camera interface package. The cameras are connected to a high-performance computer (i7-9700K, 16G RAM) equipped with a 1-terabyte SSD and 12-terabyte HDD as an image acquisition platform. The computer also controls the camera calibration module.

### SBM-VIS Dataset

The free-social behavior of two C57BL/6 mice introduced above is captured by the first version of MouseVenue3D. The first 1 min frames of four cameras are annotated as the SBM-VIS dataset, which is 7200 frames in total. To accelerate the data annotation, we take deep learning for assistance. 30% of the contours are manually labeled, and the rest are firstly labeled by YOLACT++ trained by the manually labeled 30% contours then checked by humans. Then, the single animal DeepLabCut is used to predict the poses of masked frames with the human check. Per 18 frames are grouped for a video instance and saved as YouTubeVIS format^42^. And the poses are saved as a .csv file. The identities across different cameras are corrected by human annotators.

### New scenario generation for video instance segmentation

The new scenario generation for video instance segmentation is divided into several steps: contour extraction, trajectory extraction, dataset labeling, background calculation, model self-training, and video dataset generation. After that, it can be input into the instance segmentation model for large-scale training. Suppose the number of animals in the video is n. Conda virtual environment configuration includes OpenCV 4.5.5.62, Python 3.8.12, Pytorch 1.10.1, The computer was configured with Intel(R) Xeon(R) Silver 4210R CPU @ 2.40GHz and NVIDIA RTX3090 GPU.

In the animal contour step, image thresholding is first done, and then the contour in the image is extracted, and the following formula is used to determine whether the frame is social or not, where *I* stands for a frame, *R*_i_ stands for the judgment result of this frame and *num*_*i*_ stands for the number of contours in this frame:

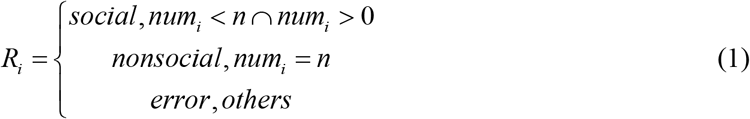

When extracting the animal trajectory, due to the influence of noise, all the contour center points are recorded as the candidates of the animal frame center point, and the closest point to each animal in the previous frame is selected from multiple center points as the true center point of this frame, and then the Hungarian matching idea is used to remove the matching points successfully, to optimize the animal trajectory.

For dataset annotation, different manually annotated datasets were used for different animals. We manually annotated 272 images in the 50 cm mice open field experiment, 805 images in the 20 cm mice open field experiment, 600 images in the birds experiment, and 800 images in the dogs experiment.

For background calculation, the non-mask position (the background) of each image is extracted and fused into the final background image using the labeled data set. The above operation is repeated for all data sets to obtain a clean background image.

The labeled data set is used for YOLACT++ round training, and the trained model is used to predict video frames. The predicted high-quality frames will be added to the original data set for the next round of training. Among them, the selection method of high-quality frames is as follows: *i* represents a certain frame, *f*_*i*_ is the segmentation result of the frame *i, f*_*i*−1_ is the segmentation result of the frame *i* −1, *F* is the calculation process of scoring matrix of all segmentation results in two frames, the calculation idea refers to the Hungarian matching idea, and the calculation result is *G*_*i*_ :

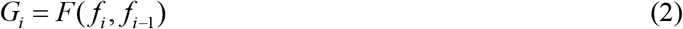

Then, all *G*_*i*_ are merged and clustered, and the class with the higher overall matrix score is selected as the high-quality frame class and added to the training data set. YOLACT++ selects the ResNet50 model as the pre-training model, and the maximum number of iterations is 150,000 generations. The training process takes about 5 hours. After YOLACT++ finishes training, its final model is used to predict the results for all frames.

The video dataset required for instance segmentation training is subsequently generated. The data set is divided into three parts, which are real data set, social area data set, and randomly generated data set. The real data set is the continuous high-quality frames predicted and filtered by YOLACT++, which are written into the video data set after data enhancement, where the data enhancement is performed by flipping the image left and right. Since there are many occlusions during social interaction and the performance of the model decreases, it is necessary to generate multiple datasets in the social area. Here, consecutive frames of animals in the social area are selected and augmented to generate the social area dataset, where *N* forms of enhancement are generated by data augmentation, as shown below, where *C* represents combination (that is, the combination of different masks is selected for flipping in each frame). *A* stands for alignment (that is, all masks are aligned to occlusion):

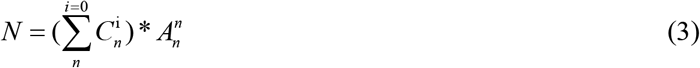

Since the number of real data and social area data sets may be far from enough to complete the model training task, some data sets in the animal activity area are randomly generated after this step. In this part, the real animal trajectory in the video, the obtained animal mask, and the background calculated in the previous step are used for data collection, and the video data set is written after data enhancement. 14940 video datasets were generated for the 50cm mice open field experiment, 15130 for the 20cm mice open field experiment, 5970 for the bird experiment, and 41,755 for the dog experiment.

### The training and validation of video instance segmentation model

Here, the video instance segmentation model adopts the Transformer-based VisTR model, which regards the video instance segmentation task as a parallel sequence encoding and decoding problem. The pre-training model was the ResNet101 model trained on the COCO dataset, the learning rate was set to 0.0001, the dropout parameter was set to 0.1, the training epochs is 30, the frame length was set to 9, the sequence length was set to 19, the number of encoding layers was 6, the number of decoding layers was 6, and Adam was used for the optimizer. The model training takes about 1.5 days. The trained model is evaluated on one minute of standard data, and the model accuracy for video instance segmentation is as follows: IST (Identity swap times) is 5.500±3.640, ISTP (Identity swap times percentage) is 0.003±0.002, IOU_NID_ (The Intersection of the union without identity) is 0.746±0.017, mAP50_NID_ (Mean of average precision without identity, the threshold value is greater than 0.5) is 0.985±0.013, mAP50_ID_ (Mean of average precision with identity, the threshold value is greater than 0.5) is 0.605±0.319, similarly, mAP70_NID_ is 0.805±0.068,mAP70_ID_ is 0.497±0.271.

### Single animal pose estimation

Single animal pose estimation was performed using DeepLabCut 2.2.0.4 with a Conda virtual environment with Python 3.8.12. Four different animals were used in the manual labeling of the dataset, with 709 images labeled for mice in a 50cm open field, 1421 images labeled for mice in a 20cm open field, 1035 images labeled for birds, and 819 images labeled for dogs. The number of body posture points varied for each animal, with 16 for each mouse(nose, left ear, right ear, neck, left front limb, right front limb, left hind limb, right hind limb, left front claw, right front claw, left hind claw, right hind claw, back, root tail, mid tail, tip tail), 19 for each bird(beak, calvaria, left eye, right eye, neck, left wing root, left wing mid, left wing tip, right wing root, right wing mid, right wing tip, left leg root, left leg tip, right leg root, right leg tip, back, belly, tail root, tail tip), and 17 for each dog(nose, left ear, right ear, neck, left front limb, left front paw, right front limb, right font paw, left hind limb, left hind paw, right hind limb, right hind paw, front back, mid back, hind back, tail root, tail tip). ResNet50 was used as the pre-trained model. The model was trained for a maximum of 103 million iterations with a batch size of 8 and took approximately 10 hours to train on an NVIDIA RTX3090 GPU using Python. The prediction results were saved in a CSV file.

### 3D pose reconstruction of multi-animals

Here, we use the multi-view geometry method in computer vision for the 3D reconstruction of multiple animals. The basic projection formula between 2D points and 3D space points is as follows.

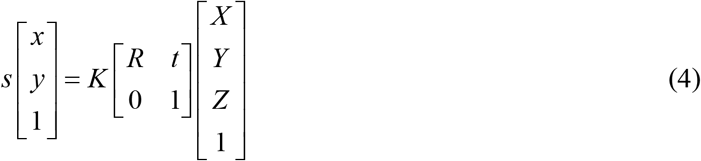

Here, *s* represents the scaling factor, *x* and *y* are the points in the image, *K* is the camera internal reference, *R* is the rotation matrix, *t* is the translation matrix, and *X, Y*, and *Z* represent the coordinates of the 3D points. Specifically, firstly, all two-dimensional skeleton information of multi-animal and multi-view was read, and the points in the two-dimensional file with too low a confidence rate were directly set to NaN. Then, the relative position parameters between multiple cameras are read and the triangulation algorithm is used for the 3D reconstruction of a single animal. The basic principle is as follows:

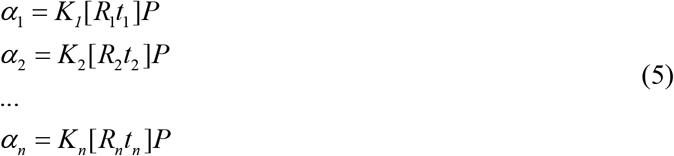

Here, *α*_1_ to *α*_*n*_ represent the two-dimensional points with the same content in different cameras, *K*_*1*_ to *K*_*n*_ represent the internal parameter matrix of different cameras, *R*_1_ to *R*_*n*_ represent the rotation matrix of different cameras, *t*_1_ to *t*_*n*_ represent the translation matrix of different cameras, and the three-dimensional point *P* can be solved by combining these equations, and we use the SVD decomposition to solve the least squares regression problem.

Next, since the appearance of animals in different views is very similar, the identities of instance segmentation may be swapped, and the wrong 3D point coordinates may be calculated. Therefore, we first obtain the full permutation index list of all 2D points of multiple animals in each view angle, and then obtain the 3D point coordinates in all cases. Eventually, the point with the smallest error is selected as the final multi-animal 3D skeleton point.

### The training and validation of animal identification model

In multi-animal experiments, because the animal hair is too similar, its identity is likely to be wrongly assigned. Here, we use the deep learning model to identify two-dimensional animals under four perspectives, to determine the identity, and to ensure that the animal identity of the whole video can be corrected back.

The data set of identity recognition needs to record the individual activity videos of all experimental animals in the same scene, and then obtain two-dimensional pictures of animals from multiple perspectives. The trained video instance segmentation model is used to predict the mask of the whole body of a single animal (the effect of manually selecting some body parts of animals for identity recognition is not good). Then, the four obtained prediction images are processed by image stretching, stitching, thresholding, and so on, and finally, a complete image is obtained as the training data. The labels are the animal numbers, so there is no need to manually annotate the data. The size of the dataset depends on the duration of recording individual activity videos of animals. In the mouse experiment with a 50 cm open field, the data set size was 594,000, in the mouse experiment with a 20 cm open field, the data set size was 180,000, in the bird social experiment, and the dog social experiment, the data set size was 54161.

The deep learning model uses the Efficientnet-b4 model, the maximum number of iterations is set to 120, the initial value of the learning rate is 0.005, and the batch size is set to 32. It is trained on NVIDIA RTX3090 GPU, and each round of training takes about 40 minutes.

In the mouse experiment with a 50 cm open field, the accuracy of the identification network in the training set was 0.993, and the accuracy of the validation set was 0.922. In the mouse experiment with a 20 cm open field, the accuracy of the training set was 0.999, and the accuracy of the validation set was 0.911. In the dog social experiment, the training set accuracy is 0.999, and the validation set accuracy is 0.999.

### The pattern visualization of animal identification by LayerCAM

LayerCAM can generate the class activation maps (CAM) of each layer of CNN-based models^25^. The LayerCAM of each layer of the EfficientNet-based identity recognition network is averaged to output a global visualization pattern of animal identities. To further compare the feature weights of different body parts of animals, the 2D poses are used for the body part location of identity frames. From the 2D poses to identity frames, there is a coordinate transformation. The transformed 2D poses on identity frames *P*_*t*_ can be calculated as:

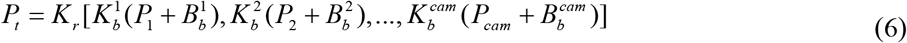

where *K*_*r*_ is the resized matrix of cascade frames, *K*_*b*_ is the scale matrix of the bounding box of single camera view, *P* is the raw 2D poses, *B*_*b*_ is the bias matrix of the bounding box of single camera view, and the index *cam* is the camera number. The *K*_*b*_ is decided by the size of frames and the bounding box size of the cropped animal instance. To reduce the disturbance of 2D pose estimation, a box centered on *P*_*t*_ of each transformed 2D pose crops the LayerCAM value. And the mean value of them represents the CAM weights of each body part.

### The mask reprojection from 3D poses to video instances

The 3D poses of each animal connect the geometric relationships of the video instances in different camera views. In the step of 3D reconstruction of multi-animals, the 2D poses of each camera view angle have been re-grouped by optimization. Because the 2D poses of multiple animals are constructed by the single animal after video instance segmentation, the masks of instances are matched to the 2D poses. Therefore, the 3D poses of each animal are corresponding to the masks of video instances frame by frame. A table saves the corresponding indexes from 3D poses to video instances and is checked frame by frame for mask reprojection.

### Parallel decomposition of trajectories

The parallel decomposition of trajectories includes three parts.

The first part is the decomposition of non-locomotor movement. Let 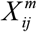 is the behavior trajectories of animals *m* with *i* frames and *j* dimensions, the non-locomotor movement component *Y*_*NM*_ can be calculated as follows:

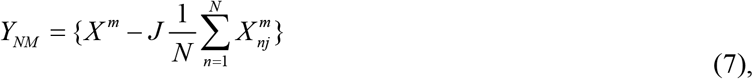

where *J* is all one vector, and *N* is the number of frames. After this step, the center of the body of the animals can be aligned together.

The second part is the decomposition of locomotion. The locomotion component *Y*_*L*_ can be .calculated as follows:

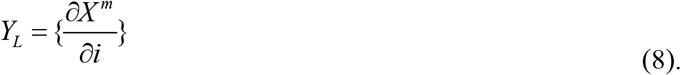

The third part is the decomposition of distance. The distance component *Y*_*D*_ can be calculated as follows:

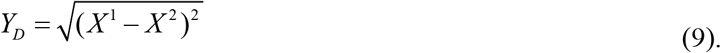

### Feature representation of distance dynamics

The distance dynamics *Y*_*DD*_ can be calculated as follows:

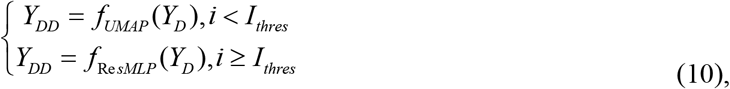

where *f*_*UMAP*_ ^(·)^ is the UMAP mapping including the parameters n_neighbors set to 199, and min_dist set to 0.3, *I*_*thres*_ is the threshold of frames set to 200000, and *f*_Re *sMLP*_ ^(·)^ is the feature representation including ResMLP. For *f*_Re *sMLP*_ ^(·)^, firstly, the *Y*_*D*_ is randomly sampled to *Y*_*Ds*_ according to *I*_*thres*_. And the rest of *Y*_*D*_ is *Y*_*Dr*_. Then, *Y*_*Ds*_ and *Y*_*DDs*_ = *f*_*UMAP*_ (*Y*_*Ds*_), the UMAP of *Y*_*Ds*_, is used to train ResMLP for feature encoding. After the training, the ResMLP predicts the *Y*_*DDr*_ from *Y*_*Dr*_, and the *Y*_*DD*_ can be recombined by *Y*_*DDs*_ and *Y*_*DDr*_ according to the sample point.

The ResMLP is based on the residual module and multi-layer perceptron^43,44^. The residual block is constructed by multi-layer perceptron with two layers. Each layer has 64 neurons, and two residual blocks are stacked to construct the residual part. The head of ResMLP is one 1d convolution layer and one global max pooling layer for the feature encoding of distance dynamics^45^. The output part of ResMLP is constructed by one fully connected layer with one sigmoid layer for the continuous value representation^46^. The activation function of ResMLP uses ReLU layers^46^. The optimizer of ResMLP is adam, the initial learning rate is set to 0.001, the mini batch size is set to 2000, and the epoch number is set to 100^47^.The final RMSE of validation is 0.02∼0.06, and the training time of ResMLP is about 4 min on NVIDIA GeForce RTX 3090 GPU.

### The time consumption comparison of ResMLP

After the manually time consumption test of UMAP, the quadratic function is used for the estimation time comparison. The coefficient of quadratic function is 0.00002. The time consumption of ResMLP is estimated as a linear function with slope set to 0.000008 and intercept set to 240 based on the training and prediction time of ResMLP. The functions of the time consumption are as follows:

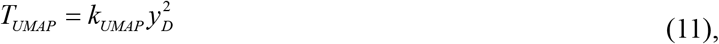

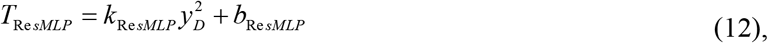

where *T*_*UMAP*_ is the time consumption of UMAP, *k*_*UMAP*_ is the coefficient of quadratic function, *y*_*D*_ is the number of distance components, *T*_Re *sMLP*_ is the time consumption of ResMLP, *k*_Re *sMLP*_ is the slope of ResMLP, and *b*_Re *sMLP*_ is the intercept.

### The distance map

Let *Y*_*E*_ is the low-dimensional embedding of the social behavior atlas, and *Y*_*DM*_ is the distance of *Y*_*E*_. The *Y*_*DM*_ can be calculated as follows:

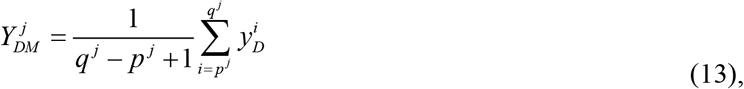

where *j* is one of the point in *Y*_*DM*_, *p* is the start time point of 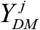, and *q* is the end time point of 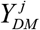.

### The map/body distance

The body distance is equivalent to *Y*_*DM*_. The map distance *Y*_*EM*_ can be calculated as follows:

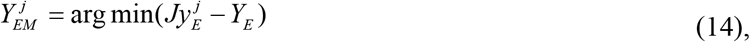

where *y*_*E*_ is one point of *Y*_*E*_. And the map/body distance *Y*_*MB*_ can be calculated as follows:

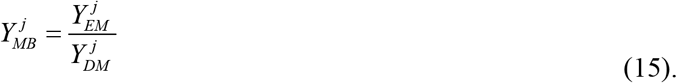

### The adaptive watershed clustering

The variable of watershed clustering on 2D embeddings is the kernel bandwidth *k*_*b*_, which decides the density *d*. The adaptive watershed clustering is designed to automatically choose the best *d*. The best *d* is determined by the stable number of clusters *c*_*st*_. To get *c*_*st*_, the clusters under certain *k*_*b*_ are firstly calculated as:

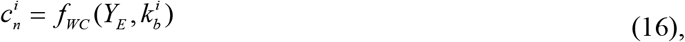

where *f*_*WC*_ ^(·)^ is the watershed clustering, *c*_*n*_ is the number of clusters. Then, the *c*_*st*_ is calculated as:

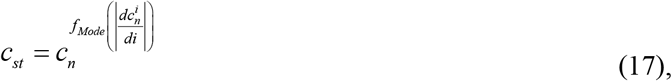

where *f*_*Mode*_ ^(·)^ is the mode function. The *c*_*s*_ is the lower bound of watershed clustering with larger kernel bandwidth. To improve the sensitivity of watershed clustering for the subtle differences of social behavior, a threshold *u*_*thres*_ is set to 0.9 to restrict *k*_*b*_ in more fine grain. So, the number of sensitivity clusters *c*_*se*_ can be calculated as:

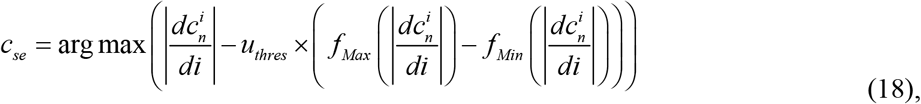

where

*f*_*Max*_ ^(·)^ is the maximum function, and *f*_*Min*_ ^(·)^ is the minimum function. The *c*_*st*_ and *c*_*se*_ together determine the lower and upper bound of watershed clustering.

### Behavior mapping of the PAIR-R24M dataset

The 3D trajectories of PAIR-R24M dataset are captured by high-performance cameras with high frame rate. To reduce the processing time and keep the global features of different mice, the frame rate is downsampled from 120 Hz to 30 Hz. The classification of the behavioral interactions of the animals includes 4 categories especially close, chase, explore and NaN value. The NaN value in social behavior atlas is defined as others. Because the interaction classes are imbalance in quantity, four coefficients are used to balance the visual effect of data distribution in atlas.

### The cluster purity

The cluster purity is an indicator to quantify the uniformity of a cluster. Let the *P* ={ *p*_1_, *p*_2_, …, *p*_*N*_ } is the ground truth indexes of all data, the *Q* = {*q*_1_, *q*_2_, …, *q*_*N*_ } is the cluster indexes of all data, and *N* is the number of clusters, the cluster purity *C*_*P*_ can be calculated as:

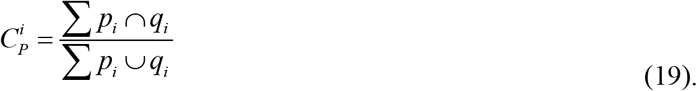

### The cluster gram of grouped mice

To reveal the inherent patterns of behavior fractions of each group, the cluster gram is firstly stacked group by group. Then, all of the behavior fractions are normalized according to the dimension of subject and sorted by hierarchical clustering according to the dimension of social behavior module. The clustering tree is normalized for better visualization. Further, the behavior fractions of each group are sorted according to Euclidean distance for the similarity metric. The initial row of each group for sorting is chose by the maximum change rate *R*_*m*_. The *R*_*m*_ can be calculated as:

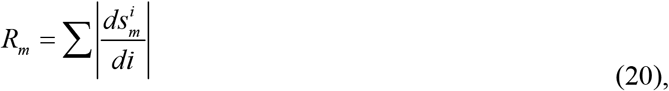

where *s*_*m*_ is the sorted social behavior fractions by hierarchical clustering.

### The angle spectrum clustering

The angle spectrum clustering is used to merge the similar sub-clusters of behavior in feature vector space. Let *V* is the feature vector matrix of social behavior modules in PCA space, the angle spectrum *A*_*s*_ can be calculated as:

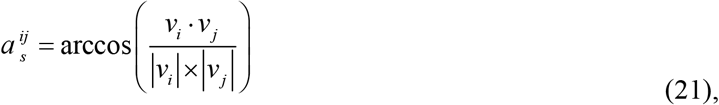

where *v* is one of the feature vector in *V*. Then, the *A*_*s*_ is clustered by hierarchical clustering according to the 11 components of 99% variance explanation.

### Computational software and hardware

The development of 3D tracking part of SBeA is based on the Python 3.8.12 in Conda environment on Ubuntu 20.04. The development of behavior mapping part and figure plot uses MATLAB R2021b. All of the statistics are finished by Prism 8.0 (GraphPad Software). The development of SBeA is on a high-performance workstation with two Intel Xeon Silver 4210R, eight NVIDIA GeForce RTX 3090, 2 Tera Byte RAM and a 140 Tera Byte Network Attached Storage. SBeA has been verified to be able to applied in a workstation with one Intel i9-12900K CPU, at least one NVIDIA GeForce RTX 3090 GPU and 128 Giga Byte RAM.

### Statistics

Before hypothesis testing, data were first tested for normality by the Shapiro–Wilk normality test and for homoscedasticity by the F test. For normally distributed data with homogeneous variances, parametric tests were used; otherwise, non-parametric tests were used. All of the ANOVA analysis are corrected by the recommended options of Prism 8.0. No data in this work are removed. All related data are included in analysis.

## Acknowledgements

This work was supported in part by STI2030-Major Projects(2021ZD0203900), National Natural Science Foundation of China (32222036), the Youth Innovation Promotion Association of the Chinese Academy of Sciences (Y2021100), the National Key R&D Program of China (2018YFA0701403), CAS Key Laboratory of Brain Connectome and Manipulation (2019DP173024), and Guangdong Provincial Key Laboratory of Brain Connectome and Behavior (2017B030301017). We thank ChatGPT for the English language editing of this paper.

## Author contributions

Conceptualization was done by YN. H., K. C. and PF. W. Code was done by YN. H., K. C., JH. L. and ZW. W. Algorithm design was done by YN. H. and K. C. Mouse data were gathered by YN. H., WH. L., XJ. W., YT. H. Bird data were gathered by YN. H., CL. H. and YT. H. Dog data were gathered by JX. L., YWZ. S., N. W., J. L., GD. W., YP. Z., YN. H., YT. H., XJ. W. and JH. L. Hardware was set up by YN. H. and K. H. Data analysis was done by YN. H. Preliminary experiments were assisted by K. H., JJ. Z., SY. C., YJ. W., G. G. and LP. W. The article was written by YN. H., K. C., YK. W. and PF. W. with input from all authors. PF. W. supervised the project.

## Supplementary materials

**Extended Data Fig. 1.**
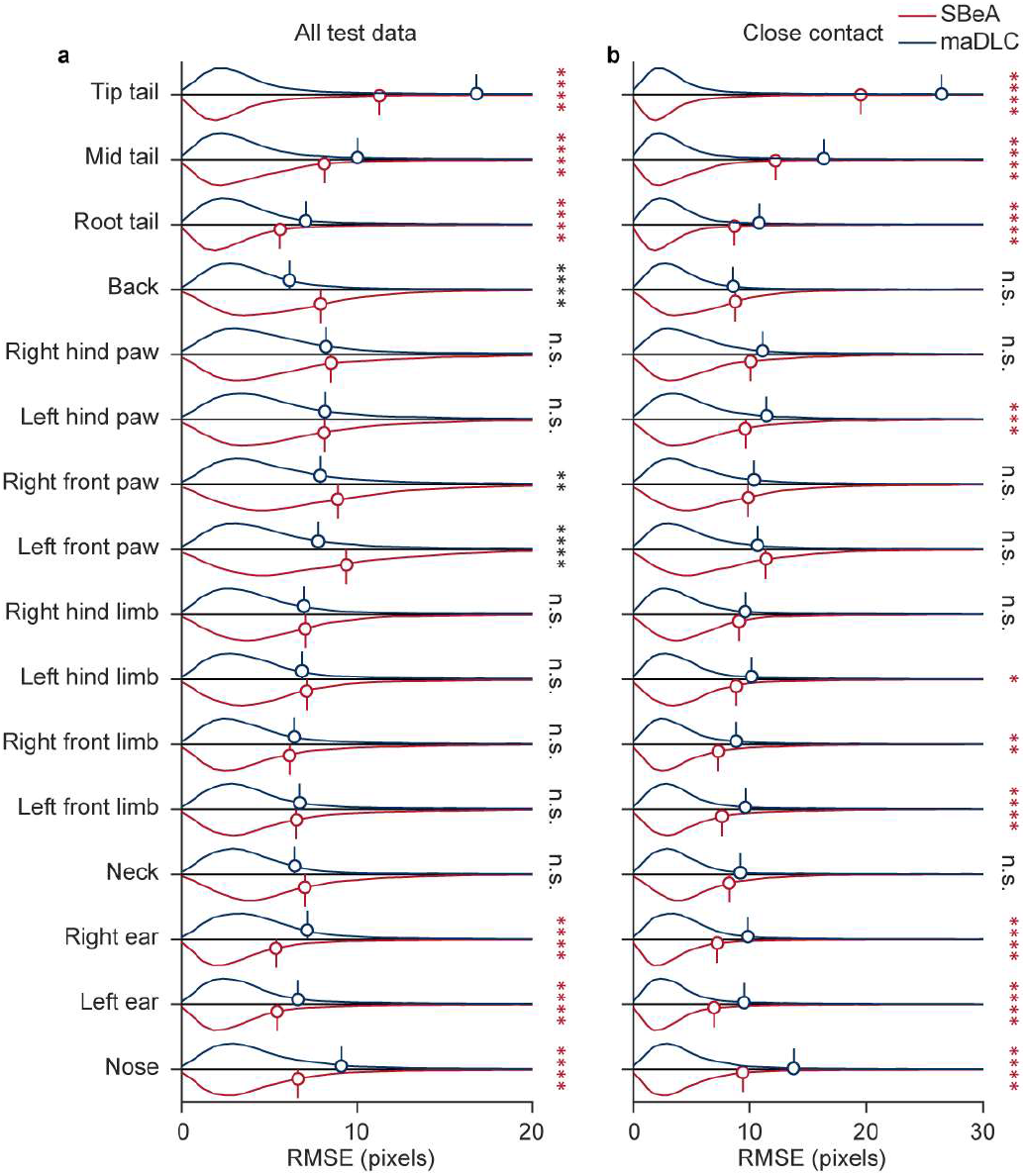
Performance comparison of SBeA and maDLC. **a**, Prediction error compasion of all test data. The RMSE of most of the body parts of SBeA is significantly lower than maDLC (two-way ANOVA followed by Sidak multiple comparisons test). **b**, Prediction error compasion of close contact. The RMSE of all of the body parts of SBeA is significantly lower than maDLC or even with maDLC (two-way ANOVA followed by Sidak multiple comparisons test). RMSE: root-mean squared error, n.s.: no significant difference, *: P<0.05, **: P<0.01, ***: P<0.001, ****: P<0.0001.

**Extended Data Tab. 1.** Social behavior definitions for manual labeling. The definition of social behavior of mouse, bird and dog refers to Mouse Ethogram database (www.mousebehavior.org), ref.^35,48–51^.

| Social behavior | Definition | Species |
| --- | --- | --- |
| Approaching | One individual approaching another individual with locomotion. | Mouse |
| Allogrooming | Grooming behavior directed towards another individual. | Mouse |
| Anogenital sniffing | Sniffing and exploring the anogenital (perianal and genital) region of another individual. | Mouse |
| Back touching | One individual touching the back of another individual by nose or fore limbs. | Mouse |
| Chasing contact | One individual running after another individual with discrete contact. | Mouse |
| Immobility | All of the individuals are motionless. | Mouse |
| Independent close grooming | All of the individuals are self-grooming without the influence of another individual. | Mouse |
| Independent rearing exploring | All of the individuals are rearing towards the outside without the influence of another individual. | Mouse |
| Locomotion rearing exploring | All of the individuals are rearing towards the outside with locomotion. | Mouse |
| Leaving back to back | All of the individuals towards back to each other and in locomotion. | Mouse |
| Micromovement | Small, subtle movements of individuals. | Mouse |
| Peer sniffing | Sniffing behavior directed towards another individual. | Mouse |
| Peer locomotion sniffing | Sniffing behavior directed towards another locomotion individual. | Mouse |
| Peer grooming sniffing | Sniffing behavior directed towards another grooming individual. | Mouse |
| Synchronous behavior | The coordinated movement or activity of multiple individuals at the same time. | Mouse |
| Anogenital touching | Contacting the genital or anal region of another individual. | Bird |
| Clamping on rectrix | One individual holding onto the tail feathers of another individual by feet. | Bird |
| Kissing | Two individuals touching beaks. | Bird |
| Back touching | One individual touching the back of another individual by nose or fore limbs. | Dog |
| Chasing | One individual pursuing or running after another individual. | Dog |
| Nose touching | Two individual touching the noses of each other. | Dog |

## References

1. Chen, P. & Hong, W. Neural Circuit Mechanisms of Social Behavior. Neuron 98, 16–30 (2018).

2. Barbera, G. et al. An open-source capacitive touch sensing device for three chamber social behavior test. MethodsX 7, 101024 (2020).

3. Sturman, O. et al. Deep learning-based behavioral analysis reaches human accuracy and is capable of outperforming commercial solutions. Neuropsychopharmacology 45, 1942–1952 (2020).

4. Schweihoff, J. F., Hsu, A. I., Schwarz, M. K. & Yttri, E. A. A-SOiD, an active learning platform for expert-guided, data efficient discovery of behavior. bioRxiv 2022.11.04.515138 (2022) doi:10.1101/2022.11.04.515138.

5. Mathis, M. W. & Mathis, A. Deep learning tools for the measurement of animal behavior in neuroscience. Curr Opin Neurobiol 60, 1–11 (2020).

6. Chen, Z. et al. AlphaTracker: A multi-animal tracking and behavioral analysis tool. bioRxiv Preprint at https://doi.org/10.1101/2020.12.04.405159 (2020).

7. Pereira, T. D. et al. SLEAP: A deep learning system for multi-animal pose tracking. Nature Methods 2022 19:4 19, 486–495 (2022).

8. Lauer, J. et al. Multi-animal pose estimation, identification and tracking with DeepLabCut. Nature Methods 2022 19:4 19, 496–504 (2022).

9. Vidal, M., Wolf, N., Rosenberg, B., Harris, B. P. & Mathis, A. Perspectives on Individual Animal Identification from Biology and Computer Vision. Integr Comp Biol 61, 900–916 (2021).

10. Agezo, S. & Berman, G. J. Tracking together: estimating social poses. Nature Methods 2022 19:4 19, 410–411 (2022).

11. Han, Y. et al. MouseVenue3D: A Markerless Three-Dimension Behavioral Tracking System for Matching Two-Photon Brain Imaging in Free-Moving Mice. Neurosci Bull 38, 303–317 (2022).

12. Huang, K. et al. A hierarchical 3D-motion learning framework for animal spontaneous behavior mapping. Nat Commun 12, (2021).

13. Han, Y., Huang, K., Chen, K., Wang, L. & Wei, P. An automatic three dimensional markerless behavioral tracking system of free-moving mice. 2021 IEEE 11th Annual International Conference on CYBER Technology in Automation, Control, and Intelligent Systems, CYBER 2021 306–310 (2021) doi:10.1109/CYBER53097.2021.9588299.

14. Liu, N. et al. Objective and comprehensive re-evaluation of anxiety-like behaviors in mice using the Behavior Atlas. Biochem Biophys Res Commun 559, 1–7 (2021).

15. Ghiasi, G. et al. Simple Copy-Paste Is a Strong Data Augmentation Method for Instance Segmentation. 2918–2928 Preprint at https://cocodataset.org/ (2021).

16. Xu, Z. et al. Continuous Copy-Paste for One-Stage Multi-Object Tracking and Segmentation. 15323–15332 Preprint at http://www.cvlibs.net/ (2021).

17. Bolya, D., Zhou, C., Xiao, F. & Lee, Y. J. YOLACT++ Better Real-Time Instance Segmentation. IEEE Trans Pattern Anal Mach Intell 44, 1108–1121 (2022).

18. Bolya, D., Fanyi, C. Z., Yong, X. & Lee, J. YOLACT Real-time Instance Segmentation. openaccess.thecvf.com https://github.com/dbolya/yolact. (2019).

19. Wang, Y. et al. End-to-End Video Instance Segmentation With Transformers. 8741–8750 Preprint at https://git.io/VisTR (2021).

20. Peça, J. et al. Shank3 mutant mice display autistic-like behaviours and striatal dysfunction. Nature 472, 437–442 (2011).

21. Mei, Y. et al. Adult restoration of Shank3 expression rescues selective autistic-like phenotypes. Nature 530, 481–484 (2016).

22. Marks, M. et al. Deep-learning-based identification, tracking, pose estimation and behaviour classification of interacting primates and mice in complex environments. Nature Machine Intelligence 2022 4:4 4, 331–340 (2022).

23. Zhuang, F. et al. A Comprehensive Survey on Transfer Learning. Proceedings of the IEEE vol. 109 Preprint at https://doi.org/10.1109/JPROC.2020.3004555 (2021).

24. Tan, M. & Le, Q. v. EfficientNet: Rethinking Model Scaling for Convolutional Neural Networks. 6105–6114 Preprint at https://proceedings.mlr.press/v97/tan19a.html (2019).

25. Jiang, P. T., Zhang, C. bin, Hou, Q., Cheng, M. M. & Wei, Y. LayerCAM: Exploring hierarchical class activation maps for localization. IEEE Transactions on Image Processing 30, 5875–5888 (2021).

26. Romero-Ferrero, F., Bergomi, M. G., Hinz, R. C., Heras, F. J. H. & de Polavieja, G. G. idtracker.ai: tracking all individuals in small or large collectives of unmarked animals. Nature Methods 2019 16:2 16, 179–182 (2019).

27. Pérez-Escudero, A., Vicente-Page, J., Hinz, R. C., Arganda, S. & de Polavieja, G. G. IdTracker: Tracking individuals in a group by automatic identification of unmarked animals. Nat Methods 11, 743–748 (2014).

28. Ebbesen, C. L. & Froemke, R. C. Body language signals for rodent social communication. Curr Opin Neurobiol 68, 91–106 (2021).

29. Bzdok, D. & Dunbar, R. I. M. The Neurobiology of Social Distance. Trends Cogn Sci 24, 717–733 (2020).

30. von Ziegler, L., Sturman, O. & Bohacek, J. Big behavior: challenges and opportunities in a new era of deep behavior profiling. Neuropsychopharmacology 1–12 (2020) doi:10.1038/s41386-020-0751-7.

31. Gomez-Marin, A., Paton, J. J., Kampff, A. R., Costa, R. M. & Mainen, Z. F. Big behavioral data: Psychology, ethology and the foundations of neuroscience. Nature Neuroscience Preprint at https://doi.org/10.1038/nn.3812 (2014).

32. McInnes, L., Healy, J. & Melville, J. UMAP: Uniform Manifold Approximation and Projection for Dimension Reduction. (2018).

33. Shi, S., Wang, Y., Dong, H., Gui, G. & Ohtsuki, T. Smartphone-Aided Human Activity Recognition Method using Residual Multi-Layer Perceptron. INFOCOM WKSHPS 2022 - IEEE Conference on Computer Communications Workshops (2022) doi:10.1109/INFOCOMWKSHPS54753.2022.9798274.

34. Wiltschko, A. B. et al. Mapping Sub-Second Structure in Mouse Behavior. Neuron 88, 1121–1135 (2015).

35. Wu, Y. E. et al. Neural control of affiliative touch in prosocial interaction. Nature 2021 599:7884 599, 262–267 (2021).

36. Marshall, J. D. et al. The PAIR-R24M Dataset for Multi-animal 3D Pose Estimation. bioRxiv 2021.11.23.469743 (2021) doi:10.1101/2021.11.23.469743.

37. Day, F. R., Ong, K. K. & Perry, J. R. B. Elucidating the genetic basis of social interaction and isolation. Nat Commun 9, (2018).

38. Wu, Y. E. & Hong, W. Neural basis of prosocial behavior. Trends Neurosci (2022) doi:10.1016/J.TINS.2022.06.008.

39. Dunn, T. W. et al. Geometric deep learning enables 3D kinematic profiling across species and environments. Nature Methods 2021 18:5 18, 564–573 (2021).

40. Mathis, A. et al. Pretraining boosts out-of-domain robustness for pose estimation. in Proceedings - 2021 IEEE Winter Conference on Applications of Computer Vision, WACV 2021 (2021). doi:10.1109/WACV48630.2021.00190.

41. Walter, T. & Couzin, I. D. Trex, a fast multi-animal tracking system with markerless identi cation, and 2d estimation of posture and visual elds. Elife 10, 1–73 (2021).

42. Yang, L., Fan, Y. & Xu, N. Video instance segmentation. in Proceedings of the IEEE International Conference on Computer Vision vols 2019-October (2019).

43. He, K., Zhang, X., Ren, S. & Sun, J. Deep residual learning for image recognition. in Proceedings of the {IEEE} conference on computer vision and pattern recognition 770–778 (2016). doi:10.1109/CVPR.2016.90.

44. Kruse, R., Mostaghim, S., Borgelt, C., Braune, C. & Steinbrecher, M. Multi-layer Perceptrons. 53–124 (2022) doi:10.1007/978-3-030-42227-1_5.

45. Kiranyaz, S. et al. 1D convolutional neural networks and applications: A survey. Mech Syst Signal Process 151, (2021).

46. Lecun, Y., Bengio, Y. & Hinton, G. Deep learning. Nature Preprint at https://doi.org/10.1038/nature14539 (2015).

47. Zhang, Z. Improved Adam Optimizer for Deep Neural Networks. in 2018 IEEE/ACM 26th International Symposium on Quality of Service, IWQoS 2018 (2019). doi:10.1109/IWQoS.2018.8624183.

48. Kort, R. et al. Shaping the oral microbiota through intimate kissing. Microbiome 2, (2014).

49. Clucas, B. Patterns of Behavior: Konrad Lorenz, Niko Tinbergen, and the Founding of Ethology. J Mammal 87, (2006).

50. Kaminski, J. & Marshall-Pescini, S. The Social Dog: Behavior and Cognition. The Social Dog: Behavior and Cognition (2014). doi:10.1016/C2012-0-06593-3.

51. de Chaumont, F. et al. Real-time analysis of the behaviour of groups of mice via a depth-sensing camera and machine learning. Nature Biomedical Engineering 2019 3:11 3, 930–942 (2019).

